# Systematic expression profiling of *dprs* and *DIPs* reveals cell surface codes in *Drosophila* larval peripheral neurons

**DOI:** 10.1101/2021.10.20.465173

**Authors:** Yupu Wang, Meike Lobb-Rabe, James Ashley, Purujit Chatterjee, Veera Anand, Hugo J. Bellen, Oguz Kanca, Robert A. Carrillo

## Abstract

In complex nervous systems, neurons must identify their correct partners to form synaptic connections. The prevailing model to ensure correct recognition posits that cell surface proteins (CSPs) in individual neurons act as identification tags. Thus, knowing what cells express which CSPs would provide insights into neural development, synaptic connectivity, and nervous system evolution. Here, we investigated expression of *dprs* and *DIPs*, two CSP subfamilies belonging to the immunoglobulin superfamily (IgSF), in *Drosophila* larval motor neurons (MNs), sensory neurons (SNs), peripheral glia and muscles using a collection of GAL4 driver lines. We found that *dprs* are more broadly expressed than *DIPs* in MNs and SNs, and each examined neuron expresses a unique combination of *dprs* and *DIPs*. Interestingly, many *dprs* and *DIPs* are not robustly expressed, but instead, are found in gradient and temporal expression patterns. Hierarchical clustering showed a similar expression pattern of *dprs* and *DIPs* in neurons from the same type and with shared synaptic partners, suggesting these CSPs may facilitate synaptic wiring. In addition, the unique expression patterns of *dprs* and *DIPs* revealed three uncharacterized MNs - MN23-Ib, MN6-Ib (A2) and MN7-Ib (A2). This study sets the stage for exploring the functions of *dprs* and *DIPs* in *Drosophila* MNs and SNs and provides genetic access to subsets of neurons.

## Introduction

During nervous system development, neurons contact thousands of cells but only form synapses with a small subset. Precise neural wiring is achieved through a series of steps including axon pathfinding, partner recognition, and synaptic pruning (Sanes and Zipursky, 2020; Zarin and Labrador, 2019). Although the mechanisms underlying these processes are not completely understood, one prevailing model proposes that cell surface proteins (CSPs) instruct chemo attraction and repulsion, self-avoidance, and synaptic partner recognition (Honig and Shapiro, 2020; Wit and Ghosh, 2016). CSPs fall into several protein families, including the immunoglobin superfamily (IgSF), the Cadherin protein family (Cdhs), the leucine-rich repeat protein family (LRRs), the receptor tyrosine kinases (RTKs), and many more (Jontes, 2017; Kurusu et al., 2008; Sanes and Zipursky, 2020; Zinn and Özkan, 2017). In vitro biochemical studies showed that subsets of these CSPs interact homo- or heterophilically, and many of these interactions are implicated in synaptic connectivity in both vertebrates and invertebrates (Cheng et al., 2019; Honig and Shapiro, 2020; Özkan et al., 2013; Wit and Ghosh, 2016).

In the well-studied vertebrate retina, retinal ganglion cells require multiple CSPs, including Sidekicks (Sdks) 1 and 2 and Dscams, to form stereotyped connections and avoid self-synapses, respectively (Garrett et al., 2018; Krishnaswamy et al., 2015; Yamagata and Sanes, 2019). Similarly, ON-OFF direction-selective ganglion cells and ON-OFF bipolar interneurons establish correct partnership by homophilic interactions of classical Cdhs (Duan et al., 2018, 2014). In hard wired invertebrate nervous systems, such as *C. elegans*, the heterophilic interaction between two IgSF proteins, Syg1 and Syg2, is required for HSNL motor neuron synapse formation (Shen et al., 2004; Shen and Bargmann, 2003). The genetically tractable *Drosophila melanogaster* is also an excellent model to study CSP expression and function due to the stereotyped neurogenesis and circuit wiring as well as extensive genetic tools. Neurons in the fly brains assemble into highly complex circuits similar to those observed in vertebrates but are numerically less daunting. In the fly mushroom body, neurons rely on different isoforms of Dscam1 to discriminate self-/non-self (Hattori et al., 2009; Wang et al., 2004; Zhan et al., 2004). In the olfactory system, epidermal growth factor (EGF)-repeat containing transmembrane Teneurin proteins, Ten-m and Ten-a, are required for the one-to-one matching between olfactory receptor neurons and projection neurons (Hong et al., 2012).

Specific challenges are also encountered in the *Drosophila* larval neuromuscular system where 33 motor neurons (MNs) within each neuromere in the ventral nerve cord (VNC) send their projections to the periphery where they follow defined paths and ultimately choose specific muscle(s) to innervate among 30 potential targets (Grueber et al., 2007; Hoang and Chiba, 2001; Menon et al., 2013). Unlike the dense neuropil where synaptic connections are difficult to identify, the neuromuscular innervation patterns are genetically hard-wired and easily identified. Thus, each efferent motor neuron can be recognized by its stereotyped innervation pattern and morphology. Utilizing the neuromuscular system, many CSPs were identified as recognition cues between MNs and muscles, including Toll (Inaki et al., 2010; Rose et al., 1997), Connectin (Nose et al., 1997, 1992) and Capricious (Kurusu et al., 2008; Shishido et al., 1998) from the LRR family and Fasciclin 2 (Davis et al., 1997; Winberg et al., 1998) and Fasciclin 3 (Chiba et al., 1995; Kose et al., 1997) from the IgSF. In contrast, the afferent neurons of the sensory nervous system are localized in the periphery and send their projections to the VNC. Forty-two sensory neurons (SNs) are stereotypically distributed throughout each hemisegment of the larval body wall and establish synaptic connections with interneurons (Orgogozo and Grueber, 2005). Studies from dendritic arborization (da) neurons identified several CSPs for self-avoidance, such as Dscam1 and Semaphorin (Meltzer et al., 2016; Miura et al., 2013; Soba et al., 2007). Thus, the unambiguous identification of cells in the motor and sensory circuits provide an ideal system to examine the genes and mechanisms that underlie synaptic specificity and development.

In a previous “interactome” screen, we and others identified two subfamilies of the Drosophila IgSF, the Defective proboscis response proteins (Dprs; 21 members) and the Dpr-interacting proteins (DIPs; 11 members) (Carrillo et al., 2015; Özkan et al., 2013). Dprs and DIPs are different from other CSPs found to wire the peripheral nervous system because they have more family members and can interact both homo- and heterophilically, providing a vast repertoire of unique combinations for synaptic specificity. Interactions between Dprs and DIPs have been implicated in synaptic connectivity, cell survival, and synaptic growth (Ashley et al., 2019; Bornstein et al., 2021; Carrillo et al., 2015; Courgeon and Desplan, 2019; Menon et al., 2019; Sanes and Zipursky, 2020; Venkatasubramanian et al., 2019; Xu et al., 2019, 2018). However, most studies focused on Dprs and DIPs have implicated only a small subset, likely due to low-penetrance targeting defects and molecular redundancy. For example, in the larval neuromuscular circuit, loss of *DIP-α* leads to complete loss of muscle 4 innervation by a specific motor neuron; however, neuromuscular junctions (NMJs) on other muscles formed by the same neuron are unaffected, suggesting different synaptic recognitions utilize different pairs of CSPs even within the same neuron (Ashley et al., 2019). Thus, obtaining a complete expression map of families of CSPs in individual neurons within specific circuits would facilitate subsequent functional studies.

Different approaches are available to map the expression patterns for genes of interest. Modern technologies like single cell RNA sequencing (scRNAseq) provide enormous information about gene expression in each cell type and has been successfully applied in the fly nervous system (Avalos et al., 2019; Li, 2020; Tang et al., 2009). However, most scRNAseq datasets do not capture the dynamic expression during development, and it is difficult to identify individual cell types from heterogenous clusters. Another approach, possibly more accurate for closely related cells, is to generate genetic reporter lines for genes of interest and directly visualize their expression. For example, in *Drosophila*, a collection of GAL4 drivers representing Gr taste receptors were used to map the projection of Gr expressing neurons (Kwon et al., 2014). These genetic reporters together with imaging allow unambiguous characterization of gene expression at a higher spatial and temporal resolution.

In this study, we interrogate the expression patterns of *dprs* and *DIPs*. These IgSF CSPs form extensive interactions and are highly enriched in the nervous system, suggesting important roles in circuit development. To access the expression of these genes, we and others generated a collection of GAL4 lines of 19 *dprs* and 11 *DIPs*. We utilized different UAS reporters to examine expression of *dprs* and *DIPs* in the *Drosophila* larval neuromuscular and sensory circuits. The distinct and stereotyped morphologies and positions of these cells allow us to unambiguously identify the reporter gene expression patterns. Here, we generated expression maps of *dprs* and *DIPs* in MNs, SNs, and muscles, and found that each MN and SN expresses a unique subset of *dprs* and *DIPs*. Utilizing hierarchical clustering, we found that the same class of SNs expresses similar *dprs* and *DIPs*, suggesting roles in identifying overlapping synaptic partners. Finally, the highly distinct expression patterns of *dprs* and *DIPs* in MNs revealed previously unidentified MNs. The expression analyses generated by this study will benefit future functional studies of Dprs and DIPs in the motor and sensory circuits. The genetic tools and pipeline provided here will facilitate expression studies of *dprs* and *DIPs*, and other CSPs, in other *Drosophila* neural circuits to promote the discovery of identification tags utilized for circuit assembly.

## Results

### Generating a GAL4 collection of *dprs* and *DIPs*

Using *Drosophila* Minos-Mediated Integration Cassette (MiMIC) insertions followed by Trojan conversion, and CRISPR-Mediated Integration Cassette (CRIMIC) insertions, we and others generated a collection of GAL4 lines of all *DIPs* and *dpr1-dpr19* (Diao et al., 2015; Kanca et al., 2019; Lee et al., 2018; Nagarkar-Jaiswal et al., 2015b; Venken et al., 2011) (Figure 1A). For each *dpr-* and *DIP-GAL4*, the cassette is inserted into a common intron or the 5’UTR shared by all isoforms (Figure 1A and Table 1). Therefore, GAL4 expression should report the expression of all isoforms of each gene. Insertion of the *SA-T2A-GAL4-PolyA* tail should generate truncated transcripts because of the presence of the PolyA tail (Logan et al., 1987; Zhang et al., 2015). In addition, the presence of a T2A-*GAL4* leads to an arrest during translation at the T2A site followed by a reinitiation of translation at the GAL4 sequence (Diao et al., 2015; Szymczak-Workman et al., 2012). To confirm the disruption of the gene of interest, we measured transcript expression by qRT-PCR using primers downstream of the insertion site and confirmed that most GAL4 lines are loss-of-function alleles. For example, in homozygous viable GAL4 lines, *DIP-α-GAL4* and *DIP-ζ-GAL4* showed no detectable *DIP-α* and *DIP-ζ* mRNA, respectively (Figure 1 – figure supplement 1A) suggesting they are null alleles. Several GAL4 lines, like *DIP-β-GAL4* and *dpr15-GAL4*, showed a reduction in mRNA levels, whereas some lines like *DIP-ι-GAL4* and *dpr16-GAL4* showed no change in mRNA expression. Although these GAL4 lines do not show a significant loss of transcription, the T2A sequence should still disrupt translation and generate mutant proteins. For homozygous lethal lines, we examined mRNA levels in heterozygous animals and found most GAL4 lines show expression near 50% (Figure 1 – figure supplementary 1B), suggesting these GAL4 lines are severe loss-of-function alleles. The qRT-PCR results are summarized in Table 2. In summary, approximately 70% of the insertions cause a severe disruption of transcription.

**Figure 1.**
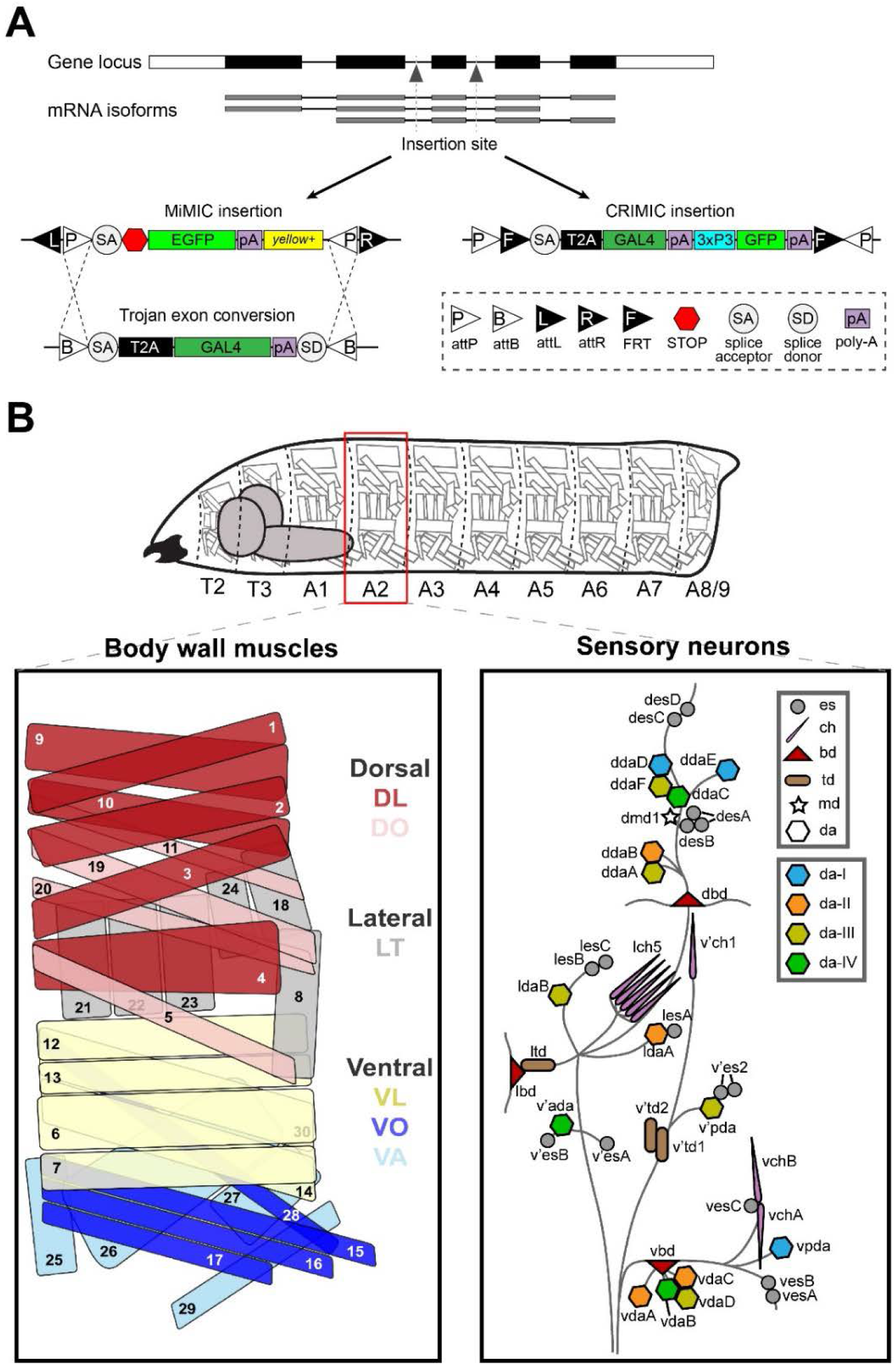
Schematic of GAL4 insertion and larval body plan. A. MiMIC or CRIMIC cassettes were inserted into a common intron or 5’UTR to capture the expression of all isoforms for each *dpr* and *DIP*. MiMIC insertions were flanked by two attP sites which are later swapped by a GAL4 exon or T2A-GAL4 trojan exon. CRIMIC insertions already carry T2A-GAL4. B. *Drosophila* larvae are divided into three thoracic segments and nine abdominal segments, with repeated muscles, MNs, and SNs. Muscles are divided into three main groups, the ventral, lateral and dorsal muscles. Ventral muscles include the ventral longitude (VL), ventral oblique (VO), ventral acute (VA) muscle groups. Dorsal muscles include the dorsal longitude (DL) and dorsal oblique (DO) muscle groups (Zarin et al., 2019). MNs innervating these muscles are not shown in this diagram. SNs are divided into six main classes: the es neurons, ch neurons, bd neurons, td neurons, md neuron, and da neurons (Orgogozo and Grueber, 2005). In addition, da neurons are further divided into da-I, da-II, da-III, and da-IV subclasses.

**Table 1.**
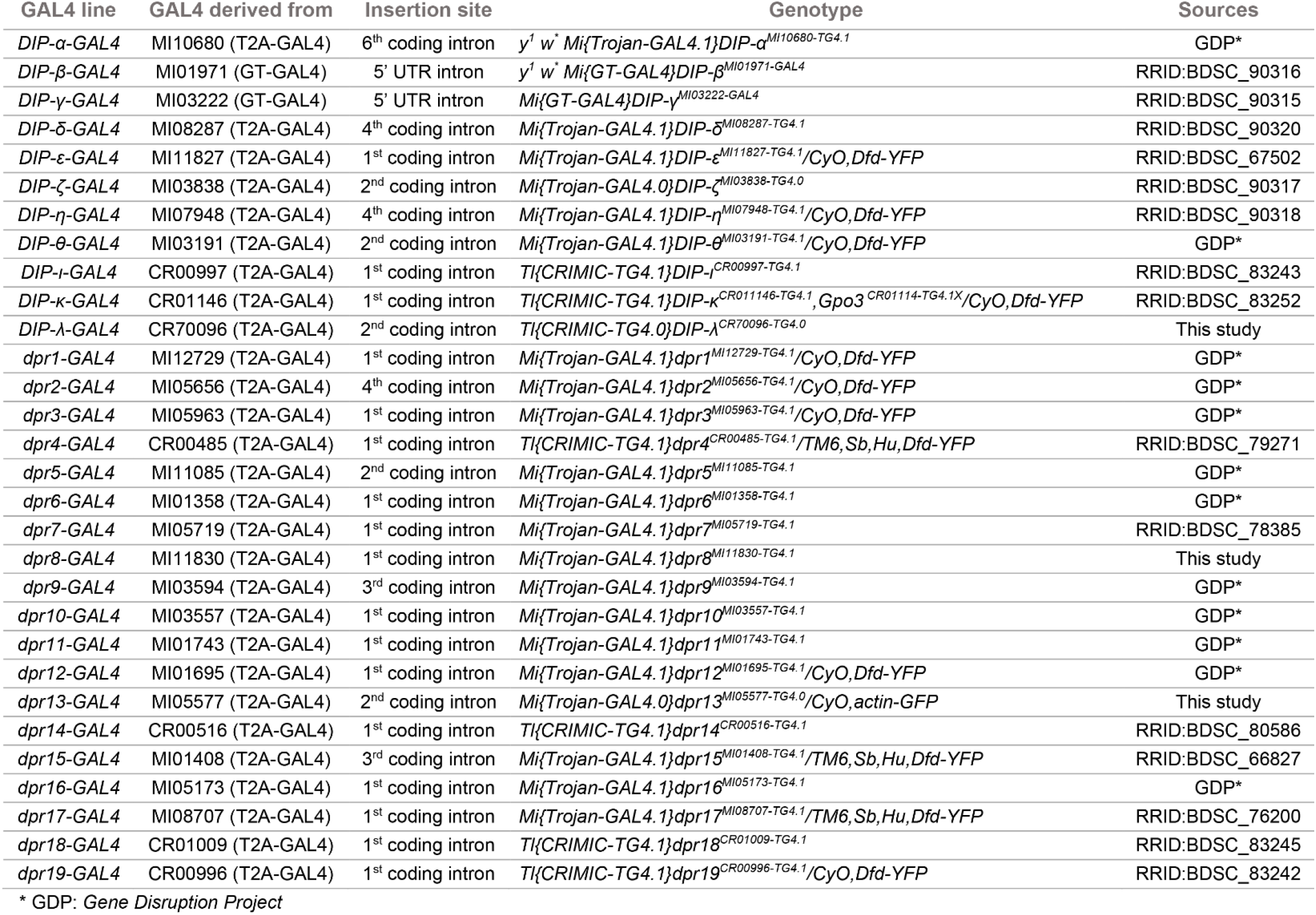

**Table 2.**
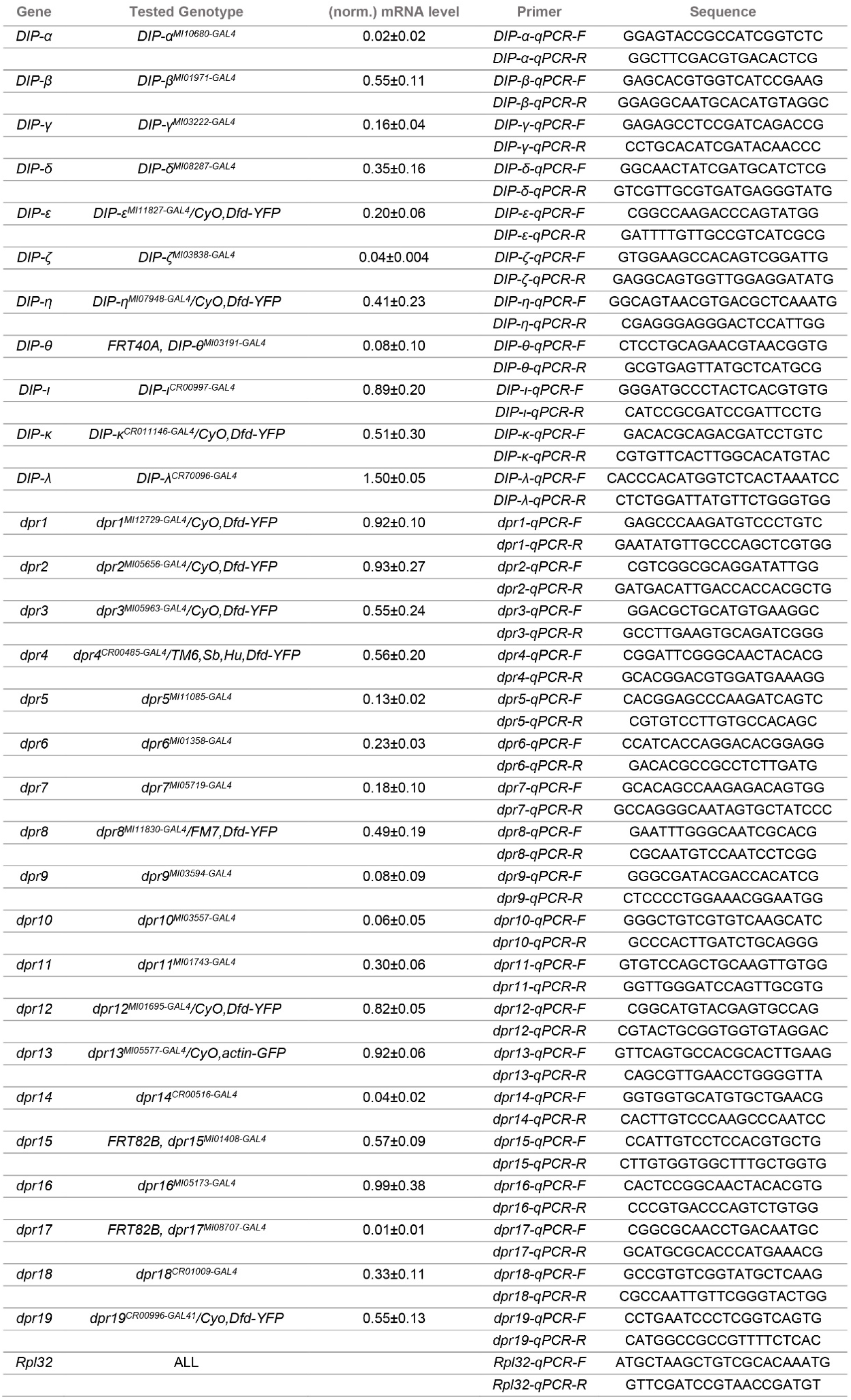

Because most GAL4 insertions are mutants, we used heterozygotes to map *dpr* and *DIP* expression. Loss of a single copy of any *dpr* or *DIP* did not affect gross viability, cell survival, or synaptic connectivity in heterozygotes as revealed by postsynaptic marker, Discs Large (DLG) and presynaptic marker, anti-horseradish peroxidase (HRP; a marker for all neuronal membranes (Jan and Jan, 1982)) (see Methods and Materials). Thus, the *dpr/DIP-GAL4* driver lines should faithfully report the cells that express *dprs* and *DIP*s (Lee et al., 2018; Nagarkar-Jaiswal et al., 2015a).

### Expression of *dprs* and *DIPs* in MNs

The larval body wall is segmented, and each abdominal hemisegment consists of 30 muscles that are grouped into three major muscle groups – ventral, lateral, and dorsal (Figure 1B) (Bate, 1990; Hooper, 1986; Zarin et al., 2019). Innervating those muscles are 33 MNs classified as type-I (29), type-II (3) and type-III (1) based on their terminal morphology and neurotransmitter type (Choi et al., 2004; Hoang and Chiba, 2001; Landgraf et al., 1997; Zarin et al., 2019). All MN axon terminals contain strings of bead-like structures called boutons which house the active zones. Type-I MNs are excitatory glutamatergic neurons, and they are further subdivided into type-I big (Ib) and type-I small (Is) due to their bouton size and innervation patterns: Ib MNs (in the larva named MN1-Ib to MN30-Ib corresponding to the muscle number) generally have larger boutons and innervate single muscle fibers whereas Is MNs have smaller boutons and innervate muscle groups (Choi et al., 2004; Lnenicka and Keshishian, 2000). The Is MN that innervates ventral muscles is referred to as the ventral common exciter (vCE), RP5, or MNISNb/d-Is, and the Is MN that innervates dorsal muscles is called the dorsal common exciter (dCE), RP2, or MNISN-Is (Broadus et al., 1995; Doe et al., 1988; Takizawa et al., 2007). Similarly, three neuromodulatory type-II MNs innervate the ventral, lateral, and dorsal muscle groups, and the single type-III MN primarily innervates m12 (Hoang and Chiba, 2001; Schmid et al., 1999). Based on these distinguishing features – terminal morphology and innervation patterns – we can unambiguously identify MNs that express each *dpr* and *DIP*.

To examine the expression of *dprs* and *DIPs* in MNs, we first crossed each GAL4 line to a fluorescent reporter line and monitored reporter expression at third instar NMJs (Figure 2A). GAL4 lines derived from MiMIC insertions were crossed to a GFP reporter, whereas CRIMIC GAL4 lines were crossed to an mCherry reporter as CRIMIC insertions carry a 3XP3-GFP marker that expresses in glial cells and the lateral bipolar dendrite (lbd) neuron (Figure 2 – figure supplement 1). To identify all NMJs, we labeled preparations with antibodies against DLG and HRP and confirmed that the gross muscle innervation was normal in *dpr/DIP-GAL4* heterozygous lines. GFP or RFP labeling of NMJs revealed the corresponding MNs that express each *dpr* and *DIP*. We followed this pipeline for each *dpr*/*DIP*-*GAL4* to record expression in all MNs.

**Figure 2.**
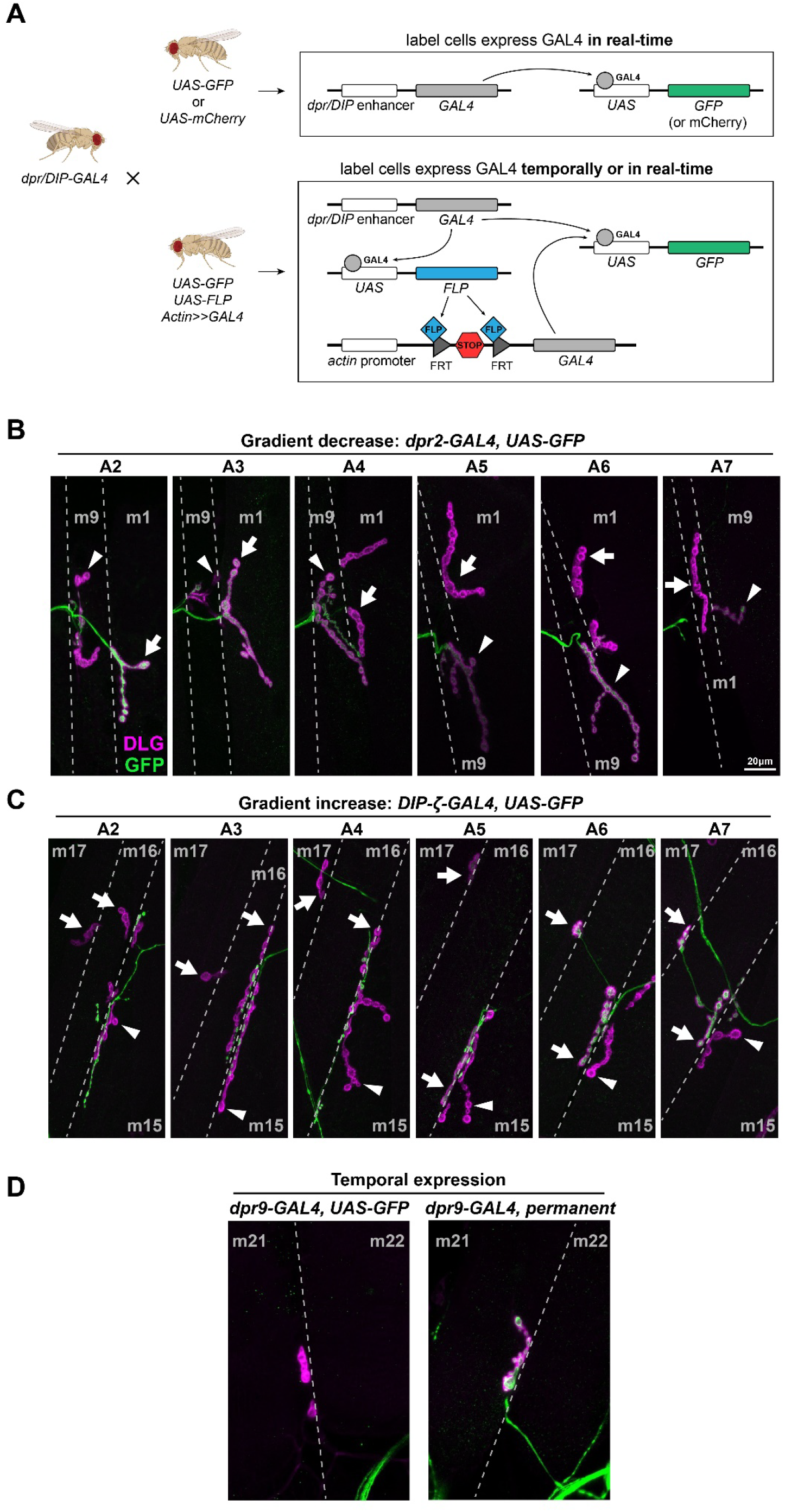
*dprs and DIPs* are expressed in various patterns in MNs. A. Schematic showing the experimental procedure. Each *dpr/DIP-GAL4* line was crossed to a real-time reporter (*UASGFP* or *UAS-mCherry*) and a permanent reporter (*UAS-GFP, UAS-FLP, actin-(FRT.STOP)-GAL4*) to reveal the dynamic expression of *dprs* and *DIPs*. B. Representative images showing an example of a decrease in expression of *dpr2-GAL4* in MN1-Ib (arrows) from anterior hemisegment A2 to posterior hemisegment A7. Note that the expression in nearby MN9-Ib (arrowheads) is also not robust as it was not expressed in A2 and A3 but expressed in A4 to A7. C. Representative images showing an example of an increase in expression of *DIP-ζ-GAL4* in MN16/17-Ib (arrows) from anterior hemisegment A2 to posterior hemisegment A7. Note that the expression in nearby MN15/16-Ib (arrowheads) was always absent. D. Representative images showing an example of temporal expression of *dpr9-GAL4* in MN21-Ib. MN21-Ib was not labeled by *dpr9-GAL4>*GFP animals, but 50% of MN21-Ib were labeled in the cross to the permanent reporter.

We mapped the expression of *dprs* and *DIPs* in all larval MNs. The expression of GAL4 and the fluorescent reporter should correlate with the endogenous gene expression. In prior work, we observed expression of *dpr6*, *dpr10*, *dpr11*, *DIP-α*, and *DIP-γ* in MNs (Ashley et al., 2019; Carrillo et al., 2015). Here, we confirmed these expression patterns; for example, *DIP-α* was selectively expressed in Is MNs but not in Ib MNs (Figure 2 – figure supplement 2A). Our data also revealed that several *dprs* and *DIPs* are not always expressed at the same level in a specific MN. For example, *DIP-δ-GAL4* only labeled 22% of abdominal MN12-Ib (Figure 2 – figure supplement 2B and 2C). Additionally, some *dprs* and *DIPs* are expressed in a gradient along the anterior to posterior axis. For example, *dpr2* showed high expression in MN1-Ib in the anterior but became undetectable from abdominal segment 4 (A4) to the posterior (Figure 2B). *DIP-ζ-GAL4*, on the other hand, labeled anterior MN16/17-Ib weakly (also known as MN15/16/17-Ib from (Hoang and Chiba, 2001; Kim et al., 2009)) but was much stronger in the posterior (Figure 2C). Note that *dpr2* also has a variable expression in MN9-Ib (Figure 2B). These complex expression patterns suggest intricate regulatory mechanisms of *dprs* and *DIPs*.

Work from our lab and others suggested that Dprs and DIPs are synaptic recognition molecules (Ashley et al., 2019; Bornstein et al., 2021; Carrillo et al., 2015; Courgeon and Desplan, 2019; Menon et al., 2019; Venkatasubramanian et al., 2019; Xu et al., 2021, 2018). In the fly neuromuscular circuit, MN axons explore the musculature field beginning in embryonic stage 14 and synaptic markers are observed in stage 16 (Yoshihara et al., 1997). A traditional UAS reporter expression in third instar larva will only report real-time expression and will not reveal if a *dpr* or *DIP* is temporally expressed earlier in development. To capture the temporal expression patterns of *dprs* and *DIPs*, we utilized a permanent labeling reporter to constantly label the GAL4-expressing neuron (Figure 2A). This method takes advantage of the FLP-out system to remove a stop codon within two FRT sites and activate an *actin*-*GAL4* to maintain GAL4 expression in any cells that expressed the gene of interest GAL4. Interestingly, we observed only a few *dprs* and *DIPs* that are temporally expressed in MNs. For example, MN21/22-Ib is not labeled when *dpr9-GAL4* is crossed to *UAS-GFP*, but with the permanent labeling reporter, the same neuron showed strong expression (Figure 2D). It is noteworthy that the CRIMIC cassettes are excisable by Flippase as well due to the presence of flanking FRT sites (Figure 1A). However, because of the activation of the permanent *actin-GAL4*, the excision of CRIMIC cassettes does not pose a technical issue.

We summarized the expression of *dprs* and *DIPs* in all MNs in Figure 3. Here, we included variable expression patterns (defined by how frequent a cell expresses the reporter) and gradient and temporal expression patterns. Criteria for each expression category is described in the Methods and in Figure 2 – figure supplement 3. In general, *dprs* are expressed in many MNs while *DIPs* are expressed much more selectively. Each MN expresses at least one *DIP*, and overall, each MN has a unique *dpr* and *DIP* expression signature. For example, we found that additional *DIPs* (*DIP-γ*, *-ε*, *-ζ*, *-η*, *-θ*, *-κ*) are also expressed in Is MNs. Interestingly, *DIP-ε* is expressed only in the ventral Is (vCE) whereas *DIP-η* is expressed only in the dorsal Is (dCE) (Figure 3). Taken together, we generated a *dpr/DIP* expression map in all larval MNs and found that each MN expresses a unique subset.

**Figure 3.**
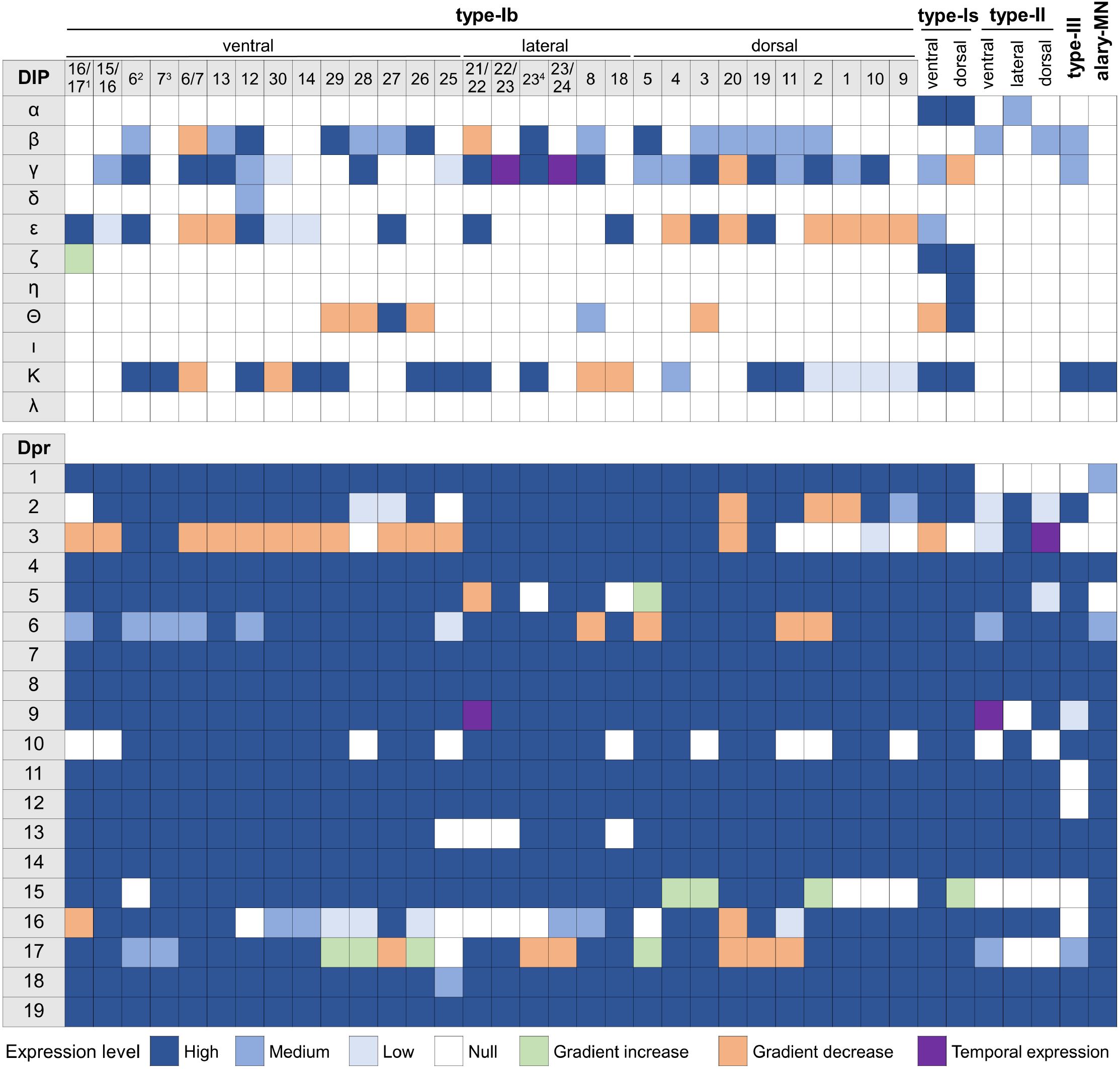
Expression map of *dprs* and *DIPs* in all larval MNs. Each column represents a MN including type-Ib, type-Is, type-II, type-III and the alary MN. Expression of each gene in each MN is characterized into a specific category as indicated in the legend. ^1^ (Hoang and Chiba, 2001) names this neuron as MN15/16/17-Ib. ^2^ Represents MN6-Ib only in A2 hemisegments, see further characterization below. ^3^ Represents MN7-Ib only in A2 hemisegments, see further characterization below. ^4^ Represents the newly identified MN23-Ib, see further characterization below.

### Expression of *dprs* and *DIPs* in SNs

Next, we examined expression of *dprs* and *DIPs* in larval SNs using similar approaches. Two morphologically distinct types of SNs can be classified in the larval body wall, and they project their axons to the VNC (Figure 1B) (Orgogozo and Grueber, 2005; Veling et al., 2019). Type-I SNs project a single dendrite that associates with chordotonal (ch) organs or external sensory (es) organs to detect mechanical and chemical stimuli. Type-II SNs are multidendritic neurons that transmit proprioceptive information. Type-II SNs can be further classified into bipolar dendrite (bd) neurons, tracheal dendrite (td) neurons, and dendritic arborization (da) neurons. The da neurons are then subdivided into four classes based on the complexity of their dendrite morphology (da-I, da-II, da-III and da-IV) (Figure 1B) (Grueber et al., 2002). SNs from the same class are uniformly distributed in the body wall into four regions: ventral, ventral’, lateral, and dorsal. The distribution of SNs enables the larva to respond to different stimuli across its entire body.

To examine expression of *dprs* and *DIPs* in SNs, we labeled larvae with anti-HRP to locate the cell bodies of SNs. The *dpr/DIP* expression map in all SNs is shown in Figure 4. Similar to MNs, *DIPs* are more sparsely expressed in SNs compared to *dprs* which are broadly expressed. However, several *dprs* (*dpr14*, *dpr15*, and *dpr17*) are only expressed in a subset of SNs, unlike their broad expression pattern in MNs. We also observed that some *dprs* and *DIPs* are temporally expressed. For example, the dorsal da neurons (ddaA, C, F and D) are labeled when *dpr5-GAL4* is crossed to the permanent labeling reporter, but not in *dpr5-GAL4>UAS-GFP* animals (Figure 4 – figure supplement 1).

**Figure 4.**
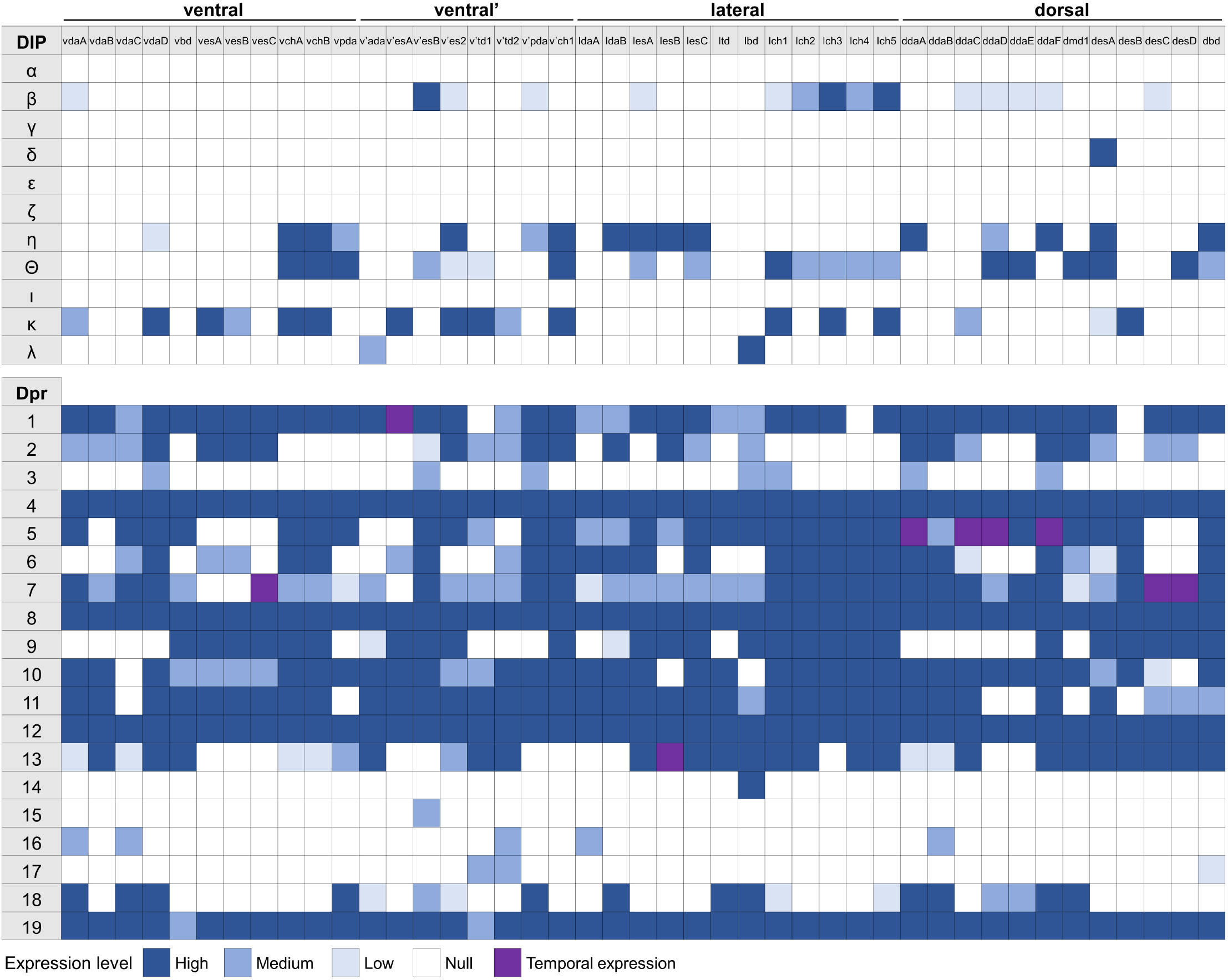
Expression map of *dprs* and *DIPs* in all larval SNs. Each column represents a SN. Expression of each gene in each SN is characterized into a specific category as indicated in the legend.

Taken together, we generated expression data for *dprs* and *DIPs* in SNs (Figure 4) and showed that each SN expresses a unique subset of *dprs* and *DIPs*, providing support for their roles as identification tags.

### SNs in the same class express similar subsets of *dprs* and *DIPs*

Larval SNs can be divided into types based on their morphology and function, including ch, es, bd, td and da neurons. Although SNs from the same type are distributed throughout the body wall and project their afferent axons through different trajectories, their axon terminals innervate the same region in the VNC and contact common interneuron partners (Grueber et al., 2007; Landgraf et al., 2003b; Merritt and Murphey, 1992; Murphey et al., 1989). For example, the ventral, ventral’, and lateral mechanosensory ch neurons project to the ventral medial region of the VNC and share synapses with several interneurons, including the Basin, Ladder, Griddle, Drunken and Even-skipped interneurons (Heckscher et al., 2015; Valdes-Aleman et al., 2021). Similarly, different classes of da neurons innervate unique sections of the VNC (Grueber et al., 2007; Merritt and Whitington, 1995; Schrader and Merritt, 2000). Overall, these innervation patterns suggest that some SNs share synaptic recognition cues while others have distinct cues.

Each SN projects a single axon into the VNC to synapse with postsynaptic targets but lacks a presynaptic neuron. Thus, SNs from the same class may share similar identification tags to wire with common interneurons. Dprs and DIPs have been implicated in synaptic partner recognition so we hypothesized that shared *dpr*/ *DIP* expression may be utilized by the same type/class of neurons to instruct synaptic specificity. To test this model, we generated an unbiased hierarchical clustering of SNs based on their *dpr/DIP* expression map (Figure 5A). Surprisingly, we found a high correlation between SN types/classes and the expression of *dprs* and *DIPs*. For example, most es neurons are grouped together, as well as all ch neurons, indicating that these two subclasses of type-I SNs can be distinguished by their expression of *dprs* and *DIPs*. Similarly, subclasses of da neurons are clustered separately. We found that da-I neurons are identifiable by expression of *DIP-θ* and the lack of *dpr2*, *dpr6*, *dpr9*, *dpr11, dpr13*, and da-II/da-III neurons are grouped by expression of *dpr2* and *dpr18,* and the lack of *dpr9* (Figure 5A). These results suggest that SNs in the same type/class may utilize similar sets of *dprs* and *DIPs* to recognize their common interneuron targets.

**Figure 5.**
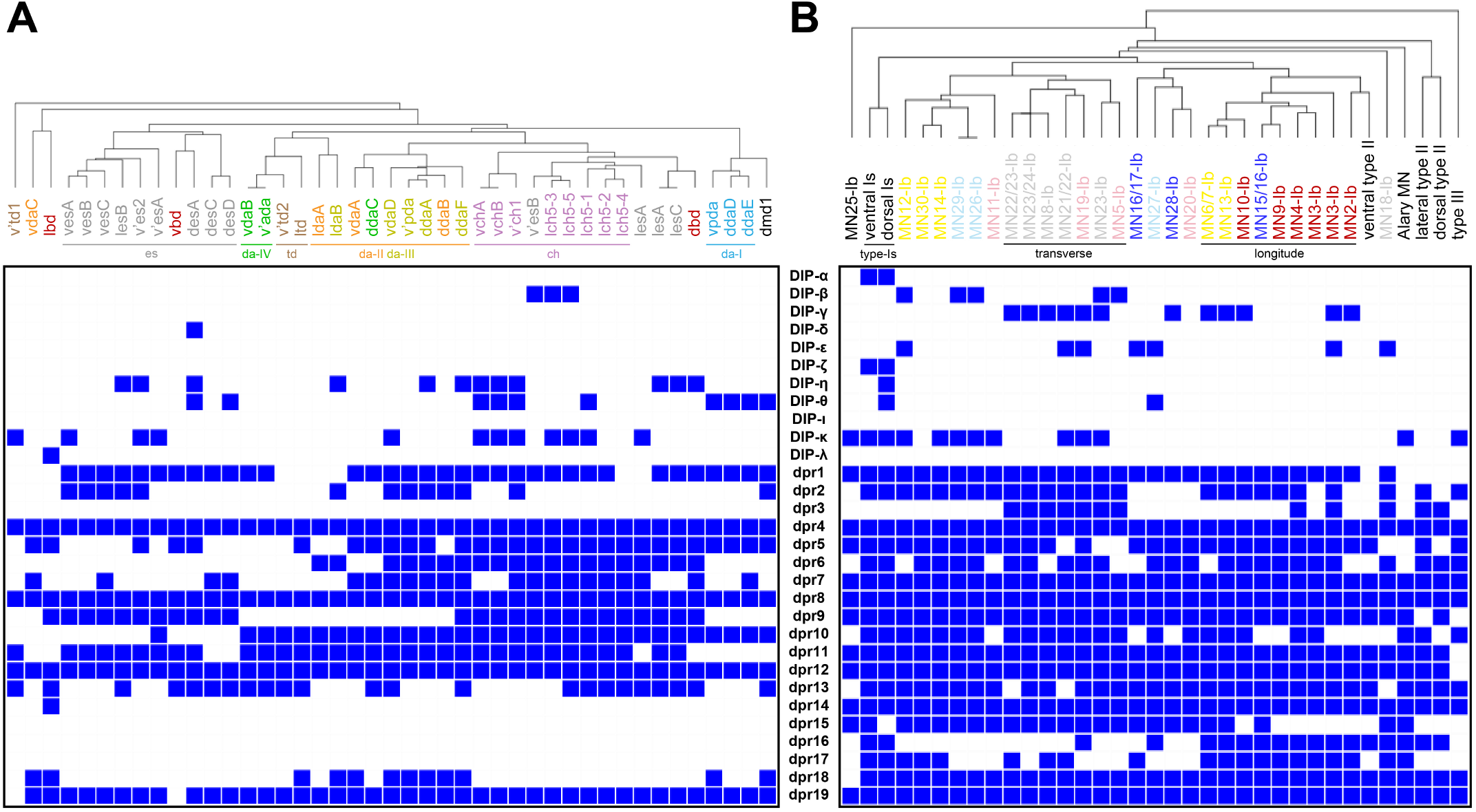
Hierarchical clustering of SNs and MNs reveals shared expression patterns of *dprs* and *DIPs* in neurons from the same class. A. SNs from the same class are clustered together based on the expression pattern of *dprs* and *DIPs*. For example, most es neurons (grey), all chordotonal neurons (purple), and da neurons fall into distinct clusters. B. Modulatory MNs (II and III) and type-Is MNs are distinct from the main type-Ib cluster. However, individual type-Ib MNs are not easily distinguished based on their expression of *dprs* and *DIPs*, except the transverse MNs (grey) and dorsal longitude MNs (red).

### *Expression of dprs* and *DIPs* is more diversified in MNs

Next, we examined if MNs that project to the same muscle groups also share the same expression patterns of *dprs* and *DIPs*. Muscles are grouped into three main spatial and functional groups – ventral, lateral, dorsal – and further divided into six subgroups based on their orientation – dorsal longitudinal (DL), dorsal oblique (DO), ventral longitudinal (VL), ventral oblique (DO) ventral acute (VA), and lateral transverse (LT) (Figure 1B) (Bate, 1990; Hooper, 1986; Zarin et al., 2019). Each muscle is normally innervated by one Ib MN and previous studies showed that Ib MNs innervating a muscle group project their dendrites to the same region in the VNC neuropil where they receive input from common premotor interneurons (PMNs) (Kim et al., 2009; Landgraf et al., 2003a, 1997; Landgraf and Thor, 2006; Mauss et al., 2009; Zarin et al., 2019). These connectivity patterns enable coordinated contraction waves that underlie larval locomotion. Thus, if Ib MNs of the same muscle group share common PMN partners, they may share similar wiring molecules. We generated an unbiased hierarchical clustering based on expression of *dprs* and *DIPs* for all MNs (Figure 5B). Type-Is, type-II and type-III MNs form independent clusters and are distinct from Ib MNs. For example, *DIP-α*, *DIP-ζ*, *dpr6* and *dpr16* are expressed in type-Is MNs, and lateral and dorsal type-II MNs are identified by the lack of *DIP-κ*, *dpr15*, *dpr17* and the expression of *dpr3* and *dpr16* (Figure 5B). However, within Ib MNs, only the MNs innervating LT and DL muscles are clustered together, whereas the other MNs appear randomly distributed. These results suggest that based on the expression patterns of *dprs* and *DIPs*, MNs can be clustered by their type, but Ib MNs cannot be further clustered by the muscles they innervate. MNs must identify not only their presynaptic inputs, but also their distinct postsynaptic partners. The combination of pre- and postsynaptic partnerships may explain the inability to cluster Ib MNs based on their expression patterns of *dprs* and *DIPs*. Therefore, more complex identification codes may be necessary for MNs to distinguish both pre- and postsynaptic partners.

### *dpr*/*DIP* expression maps reveal additional MNs

#### Alary muscle MN

In addition to the muscles required for larval locomotion, larvae have another segmentally repeated muscle – the alary muscle – that attaches to the trachea along the larval heart tube (Bataillé et al., 2015). Although the morphological and functional properties of the alary muscle have been examined (Boukhatmi et al., 2014), the development, connectivity, and functional properties of the MN that innervates alary muscles are still lacking. The alary muscle MN axon resides in the transverse nerve (TN) and projects along m8 towards the alary muscle. Here, we mapped the expression of *dprs* and *DIPs* in the alary muscle MN. As previously observed, the dendrite of the lbd neuron travels in parallel with the alary muscle MN axon within the TN (Gorczyca et al., 1994; Macleod et al., 2003; Thor and Thomas, 1997). Thus, if a *dpr* or *DIP* is expressed in both the alary muscle MN and lbd, we would be unable to distinguish them in the nerve. Therefore, we monitored the co-localization of DLG and the fluorescent reporter on the alary muscle to unambiguously assign expression (Figure 3 – figure supplement 1). We observed that alary muscle MN NMJs share features of type-I boutons including the size and DLG labeling surrounding the boutons (type-I boutons are surrounded by significant DLG (Guan et al., 1996)). Using the same criteria described for MNs and SNs, we found one *DIP* and many *dprs* that are expressed in the alary muscle MN, including *DIP-κ*, *dpr4*, and *dprs7-19*. These expression data and driver lines will facilitate future characterization of this MN.

#### MN23-Ib

Most Ib MNs have a single muscle target. However, some Ib MNs innervate two muscles in close proximity, likely due to shared recognition cues. For example, a previous study found that Ib MNs innervating the lateral muscles can synapse with neighboring muscles and thus named these neurons MN21/22-Ib, MN22/23-Ib, and MN23/24-Ib (Figure 6A) (Hoang and Chiba, 2001). These innervation patterns were later confirmed by MARCM analysis (Kim et al., 2009).

**Figure 6.**
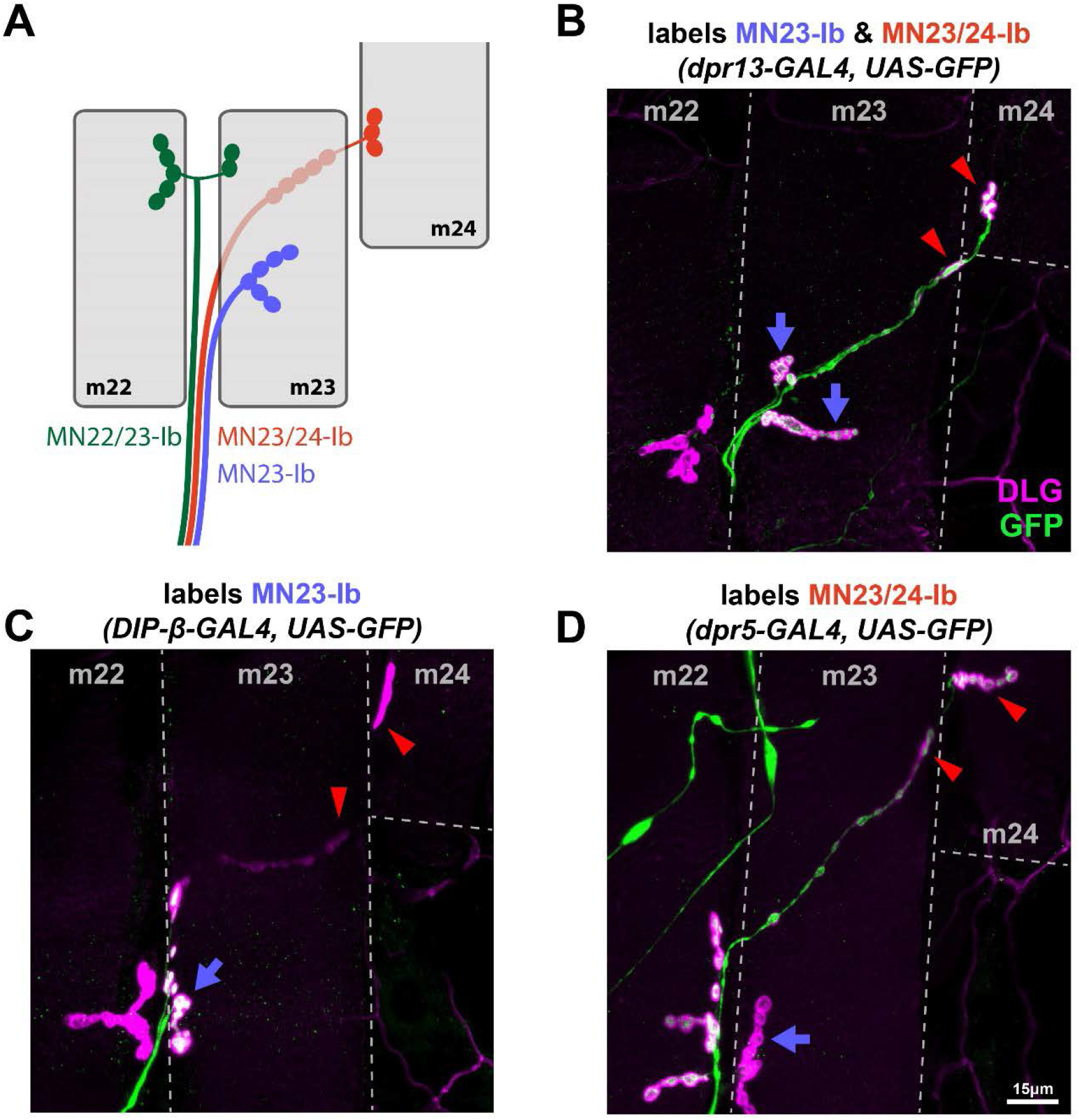
Differentially expressed *dprs* and *DIPs* reveal a MN that solely innervates m23. A. Schematic depiction of transverse muscles 22, 23 and 24 (grey) with previously identified MN22/23-Ib (green), MN23/24-Ib (red) and newly identified MN23-Ib (blue). MN22/23-Ib innervates the cleft between m22 and m23. MN23/24-Ib travels underneath m23 and forms boutons on m23 and m24. MN23-Ib only innervates m23. B. Representative image showing *dpr13-GAL4* expression in both MN23/24-Ib (red arrowheads) and MN23-Ib (blue arrows). Thus, all boutons on m23 and m24 are labeled by GFP. C. Representative image showing *DIP-β-GAL4* expression in MN23-Ib (blue arrow). Boutons underneath m23 and boutons from m22, m24 (red arrowheads) are not labeled by GFP, thus *DIP-β-GAL4* is not expressed in MN22/23-Ib and MN23/24-Ib. D. Representative image showing *dpr5-GAL4* expression in MN22/23-Ib and MN23/24-Ib (red arrowheads), but not in MN23-Ib (blue arrow). The lack of GFP in the arbor on m23 indicated the existence of a MN that solely innervates m23.

In our analyses, we observed that m23 has several Ib NMJ branches and m24 has only one NMJ (Figure 6B). While a single MN can form several branches on a muscle, we found some *dpr*/*DIP-GAL4s* that only label one Ib branch on m23 and no other branches on lateral muscles (Figure 6C). These data suggest the existence of an additional MN that solely innervates m23, and we named it MN23-Ib. The bouton size and DLG labeling intensity of M23-Ib boutons indicates that it is a type-Ib NMJ. *DIP-β* and *DIP-κ* are expressed in MN23-Ib and not in the nearby MN22/23-Ib and MN23/24-Ib (Figure 6C). Note that MN23/24-Ib forms long, linear Ib NMJs on the underside of m23 before it reaches m24 (Figure 6A and 6C). We also found that *dpr5* was expressed in MN23/24-Ib and nearby MN22/23-Ib, but not in MN23-Ib (Figure 6D), providing further evidence for an additional Ib MN solely innervating m23. Additional *dprs* and *DIPs* are expressed in both MN23-Ib and MN23/24-Ib (Figure 6B). Thus, we describe a previously unidentified Ib MN that innervates m23.

#### MN6-Ib and MN7-Ib in A2

Another example of a dual-targeting Ib MN is MN6/7-Ib (also known as RP3 in the embryo) (Schmid et al., 1999; Sink and Whitington, 1991a, 1991b). The innervation pattern of MN6/7-Ib was initially identified by dye fill labeling and MARCM (Hoang and Chiba, 2001; Kim et al., 2009; Sink and Whitington, 1991b). Due to the ease of accessibility of m6 and m7, MN6/7-Ib is extensively used for studies of synaptic connectivity, synaptic growth, and synaptic homeostasis.

Based on these previous studies, we predicted that if a *dpr* or *DIP* were expressed in MN6/7-Ib, the Ib NMJs on both m6 and m7 would be completely fluorescently labeled (Figure 7A, right). Surprisingly, in A2, we observed several *dpr*/*DIP-GAL4s* that are expressed in Ib MNs that have large NMJs on m6 and others that are expressed mainly in the Ib NMJs on m7 (Figure 7A, left). For example, *DIP-β, DIP-γ* and *DIP-ε* were expressed in a MN that mainly innervates m6 (Figure 7B), whereas *dpr15* was expressed in a MN that mainly innervates m7 (Figure 7C). These expression patterns suggested that two Ib MNs innervate m6 and m7 in A2. Hereafter, we named these MNs as MN6-Ib and MN7-Ib. Prior studies hinted at the possibility of two MNs based on the larger synaptic terminal area on m6/7 in A2 compared to A3-A6 (Lnenicka and Keshishian, 2000). However, it was thought that the larger synaptic area was due to a large NMJ from a single Ib MN. In the larval neuromuscular circuit, the number of boutons reflect the size of the NMJ. We quantified the m6 and m7 Ib NMJs and observed a significantly larger arbor in A2 compared to A3 (Ib NMJ on m6: 34.2 on A2 and 18.5 on A3; Ib on m7: 23.1 on A2 and 11.7 on A3) (Figure 7 – figure supplement 1). Taken together, m6 and m7 in A2 are innervated by two Ib MNs.

**Figure 7.**
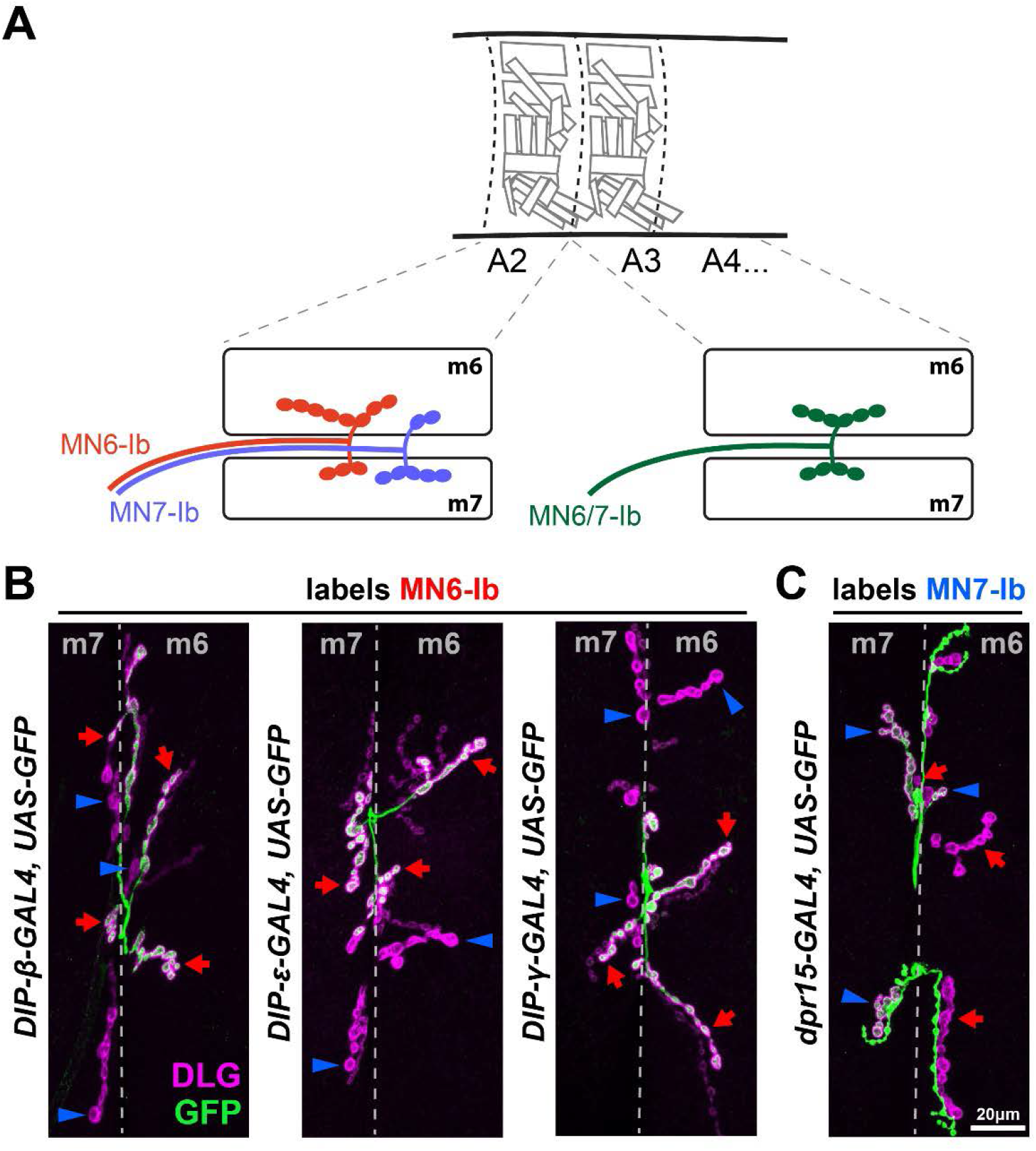
Differentially expressed *dprs* and *DIPs* reveal MN6-Ib and MN7-Ib in segment A2. A. Schematic depiction of MN6-Ib (red) and MN7-Ib (blue) in segment A2, and MN6/7-Ib in A3-A7 (green). MN6-Ib preferentially innervates m6 but also forms a small NMJ on m7, whereas MN7-Ib prefers m7 but also forms a small NMJ on m6. B. Representative images showing that *DIP-β*, *DIP-ε* and *DIP-γ* are specifically expressed in MN6-Ib (red arrows), but not in MN7-Ib (blue arrowheads). Note that MN6-Ib forms boutons with both m6 and m7, since there is a small GFP positive type-Ib NMJ on m7 (red arrows on m7). Conversely, the lack of GFP in most m7 type-Ib NMJ and the small m6 type-Ib NMJ (blue arrowheads) indicate MN7-Ib also dual innervates both muscles. C. Representative image showing that *dpr15* is specifically expressed in MN7-Ib (blue arrows) but not in MN6-Ib (red arrowheads). MN6-Ib and MN7-Ib also show dual innervation patterns in this genetic background.

### Characterization of MN6-Ib and MN7-Ib

A recent study reported a GAL4 driver (*GMR79H07-GAL4*) that labels MN6-Ib in A2 (Aponte-Santiago et al., 2020). We tested this driver and confirmed MN6-Ib expression; however, it sometimes labels MN7-Ib NMJs or both MN6-Ib and MN7-Ib, suggesting that this reporter is not specific to MN6-Ib (Figure 8 – figure supplement 1). We examined MN6-Ib and MN7-Ib further to better understand their innervation patterns and dendritic projections. Interestingly, we found that MN6-Ib and MN7-Ib preferentially innervate their corresponding muscle, but sometimes, these MNs also form minor NMJs on the neighboring muscle (Figure 7A). Next, we monitored the frequency of dual innervation of each MN using *GMR79H07-GAL4* and found that 68.2% of MN6-Ib and 72.7% of MN7-Ib innervate both muscles (Figure 8A). We also determined the size of each NMJ by counting Ib boutons and found that on average MN6-Ib forms 48.6 boutons on m6 and 5.9 boutons on m7, while MN7-Ib forms 3.1 and 30.8 boutons on m6 and m7, respectively (Figure 8B).

**Figure 8.**
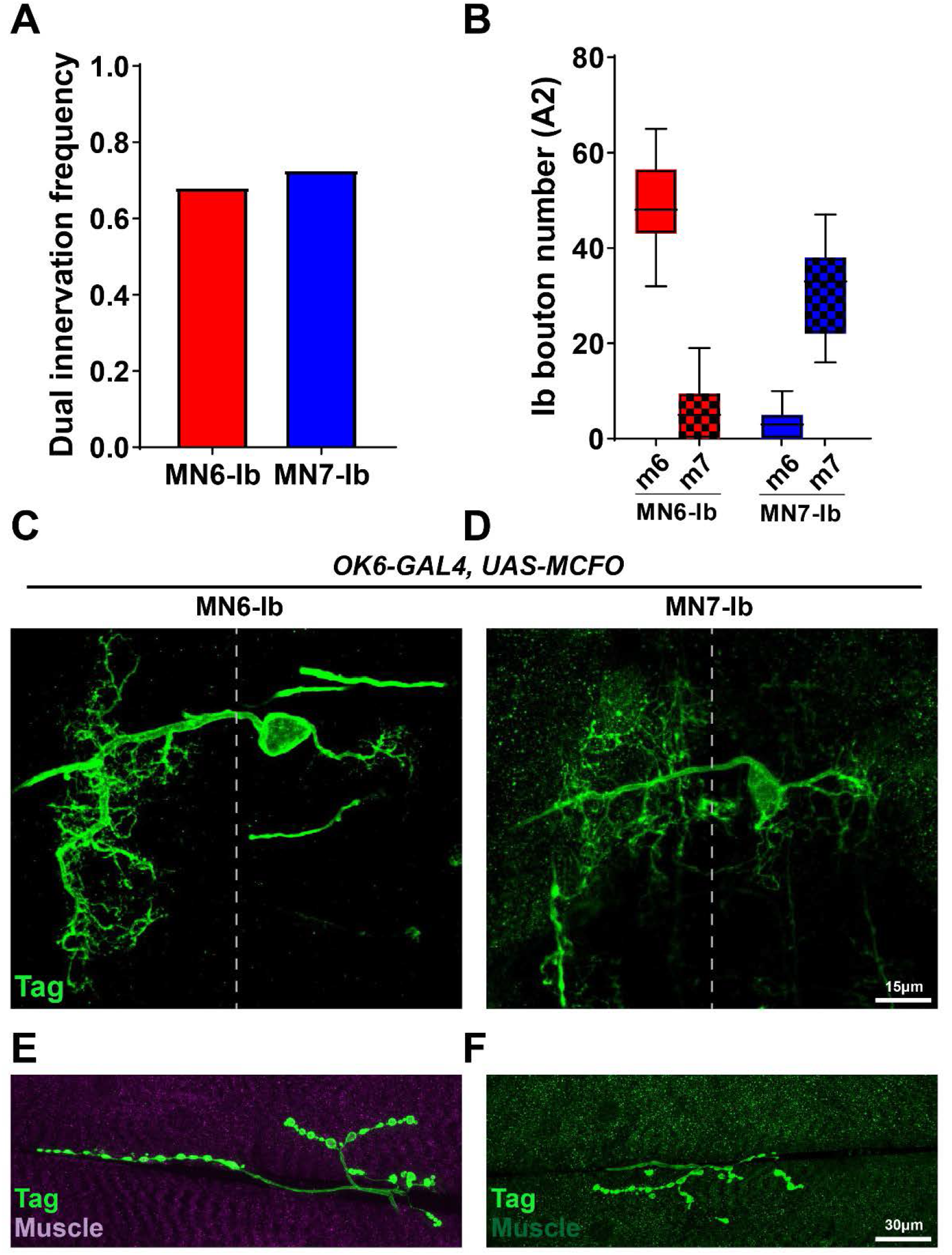
Further characterization of MN6-Ib and MN7-Ib. A. Quantification of the dual innervation frequencies of MN6-Ib and MN7-Ib. 68.2% of MN6-Ib also innervate m7 and 72.7% of MN7-Ib also innervate m6. B. Quantification of MN6-Ib and MN7-Ib NMJ sizes on both muscles. On average, MN6-Ib forms ∼49 boutons with m6 and ∼6 boutons with m7, while MN7-Ib forms ∼3 boutons with m6 and ∼31 boutons with m7. C-D. A pan MN driver *OK6-GAL4* driving MCFO revealed the dendritic morphology of MN6-Ib and MN7-Ib in the VNC. Both MNs have similar morphologies including large contralateral dendritic arbors and small ipsilateral arbors. E-F. Corresponding NMJ images from the same neuron shown in C (MN6-Ib) and D (MN7-Ib)

MN6/7-Ib (RP3) is derived from neuroblast 3-1 (NB3-1) (Schmid et al., 1999; Sink and Whitington, 1991a, 1991b). To visualize the dendritic projections of MN6-Ib and MN7-Ib in A2, we examined their cell body position in the VNC and dendrite morphology using a pan MN driver, *OK6-GAL4*, with multi-color FLP-out (MCFO) (Nern et al., 2015). We found that the cell bodies of MN6-Ib and MN7-Ib are both localized at the dorsal neuropil and project axons to the contralateral hemisegment. They also extend a small dendritic arbor to the ipsilateral side (Figure 8C-F). These features are shared with RP3 (MN6/7-Ib) (Kim et al., 2009). These data suggest that these two MNs likely both originated from NB3-1. Overall, we identified and confirmed the presence of two Ib MNs in A2 that preferentially innervate m6 or m7.

### Expression of *dprs* and *DIPs* in the glial cells

In the Drosophila larval peripheral nervous system, glia plays important roles in neuronal development, axon path finding, and synaptic homeostasis (Bittern et al., 2020; Yildirim et al., 2019). In segmental nerves, sub-perineural glia and wrapping glia form extensive interactions with the axons (Kottmeier et al., 2020). Therefore, we examined the glial expression of *dprs* and *DIPs*.

In the previous analyses using UAS-GFP and the permanent labeling reporter, we were unable to unambiguously distinguish glial and neuronal expression due to labeling of the MN axons in each segmental nerve. Therefore, to probe glial expression, we crossed each *dpr/DIP-GAL4* line with a nuclear reporter line, G-TRACE (Evans et al., 2009). The G-TRACE system utilizes FLP-FRT and GAL4-UAS to report both temporal and real-time gene expression (Figure 9A). If a GAL4 is transiently expressed, then cell nuclei will be GFP positive. However, if the cell nuclei are labeled by both GFP and RFP, this may suggest the GAL4 is consistently expressed. We crossed each *dpr/DIP-GAL4* line to the G-TRACE reporter and found that *dpr1* is expressed in glia (Figure 9 -figure supplement 1). Additionally, *dpr1* expression is highly dynamic since some glia temporarily express *dpr1* while others maintain *dpr1* expression. Overall, *dpr1* was only expressed in a subset of glia, and the restricted expression of *dprs* and *DIPs* suggest that these CSP subfamilies have limited roles in glial cells.

**Figure 9.**
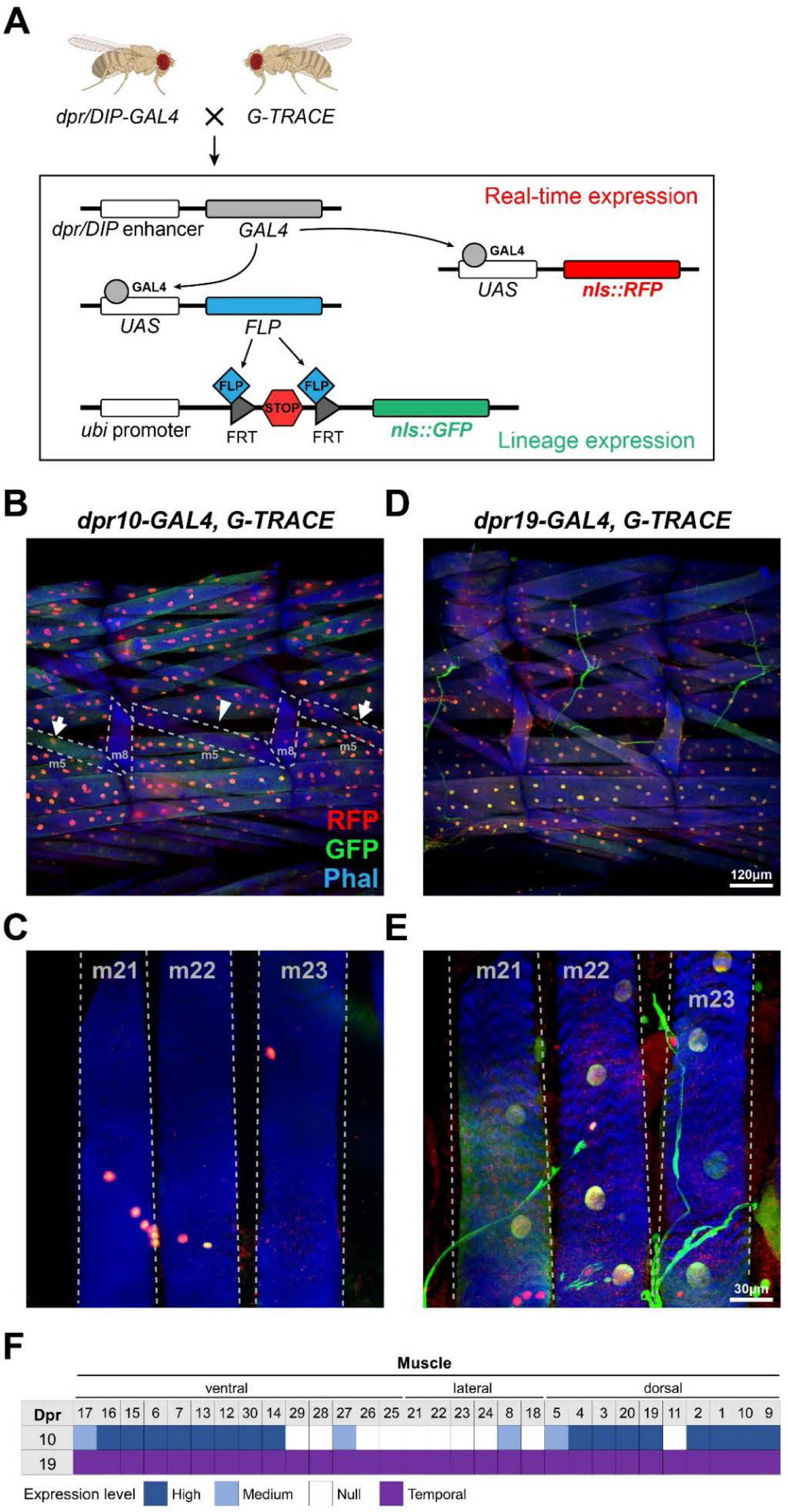
Using the GTRACE system to probe expression of *dprs* and *DIPs* in muscles and glial cells. A. Schematic depiction showing the cross between *dpr/DIP-GAL4* and the *GTRACE* reporter. Red signal represents real-time GAL4 expression and green signal represents earlier GAL4 expression. B-C. *dpr10* is consistently expressed in most muscles (B) but absent in transverse muscles (C) and some deeper ventral muscles. Expression in some muscles is not consistent. For example, in some hemisegments m5 nuclei are not labeled (arrowhead), but an adjacent hemisegment shows labeling of m5 nuclei (arrows). *dpr10* expression is maintained throughout development as revealed by co-labeling with GFP and RFP. D-E. *dpr19* is expressed in all muscles (D) including transverse muscles (E). Compared to *dpr10*, these nuclei have less RFP intensity, which may indicate that *dpr19* is temporally expressed in early development and turned off later. F. Expression map of *dpr10* and *dpr19* in muscles.

### Expression of *dprs* and *DIPs* in muscles

In a previous study, we observed *dpr10* expression in ventral and dorsal muscles and its interacting partner, *DIP-α*, in Is MNs (Ashley et al., 2019). Loss of Dpr10-DIP-α interactions cause a complete loss of Is MN innervation on m4 while other Is NMJs were unaffected, suggesting that other pairs of synaptic recognition molecules are involved in Is MN-muscle recognition (Ashley et al., 2019). To examine the role of other *dprs* and *DIPs* in this process, we mapped their expression in muscles utilizing the G-TRACE system to label muscle nuclei. We first confirmed expression of *dpr10* in all longitudinal muscles, but not in oblique or transverse muscles (Figure 9B,C). This expression pattern suggests distinct transcriptional regulation programs between muscle groups (Bate, 1990). In some hemisegments, a small subset of muscles, such as m5 and m8, showed inconsistent expression of *dpr10* (Figure 9B, arrow). Also, all muscle nuclei co-labeled with GFP and RFP in *dpr10-GAL4>G-TRACE* (Figure 9B), suggesting that *dpr10* expression is maintained throughout larval development.

We examined other *dprs* and *DIPs* and found that *dpr19* is expressed in all muscles (Figure 9D), including the oblique and transverse muscles (Figure 9E). Unlike *dpr10*, most muscle nuclei in *dpr19-GAL4>G-TRACE* are only GFP positive, suggesting that *dpr19* is temporally expressed and turned off in late larval stages. To support this temporal expression, we examined *dpr19-GAL4*>*mCherry* first instar larva and observed high level of muscle expression, which we did not observe in third instar (Figure 9 – figure supplement 2). Taken together, we showed that muscles express many fewer *dprs* and *DIPs* compared to motor and sensory neurons (Figure 9F). These results suggest that a subset of *dprs* and *DIPs* may function in the MN-muscle recognition and others in PMN-MN recognition.

## Discussion

Dprs and DIPs play important roles in nervous system development. To date, only a small subset of Dpr-DIP pairs has been examined, including Dpr11-DIP-γ, Dpr6/10-DIP-α, and Dpr12-DIP-δ. These CSP pairs were implicated in synaptic recognition (Ashley et al., 2019; Bornstein et al., 2021; Venkatasubramanian et al., 2019; Xu et al., 2019, 2018), neuronal survival (Courgeon and Desplan, 2019; Menon et al., 2019; Xu et al., 2021), and synaptic growth (Carrillo et al., 2015). Furthermore, Dprs and DIPs are widely expressed across many neural circuits. Several labs have utilized the GAL4/UAS system to visualize expression of *dprs* and *DIP*s in olfactory neurons (Barish et al., 2018), adult leg MNs and SNs (Venkatasubramanian et al., 2019), optic lobe neurons (Cosmanescu et al., 2018), and fru P1 neurons (Brovero et al., 2021). While these studies revealed unique *dpr*/*DIP* expression in the respective neurons, the depth of the expression map was limited due to the less complete GAL4 collection at the time, and some studies only focused on a global expression pattern without characterization of individual cell types.

Here, we reported a collection of GAL4 enhancer trap lines for all *DIPs* and 19 *dprs*, and examined their expression in larval MNs, SNs, peripheral glia, and muscles. Interestingly, each neuron expresses a unique combination of *dprs* and *DIPs*. We also found that many *dprs* and *DIPs* are expressed in patterns including different expression levels, in anterior-posterior gradients, and temporal expression. Surprisingly, our expression analysis also revealed previously uncharacterized larval MNs that differentially express *dprs* and *DIPs*. Finally, we showed that *dpr10* and *dpr19* are expressed in muscles, suggesting that additional Dpr-DIP interactions may instruct MN-muscle recognition. The *dpr*/*DIP* expression map identified here, along with the GAL4 lines that are also hypomorphs or loss-of-function alleles, will facilitate examination of Dpr-DIP interactions in development of motor, sensory, and many other circuits.

### Using the *dpr/DIP* code to annotate single cell RNA sequencing data

Recent advances in single cell RNA sequencing (scRNAseq) provide a powerful, high-throughput approach to identify large scale gene expression patterns. Various *Drosophila* neural tissues have been analyzed by scRNAseq, including the adult brain (Davie et al., 2018), optic lobe (Konstantinides et al., 2018), adult VNC (Allen et al., 2020; Genovese et al., 2019), larval brain (Avalos et al., 2019), eye disc (Ariss et al., 2018) and the larval VNC (Nguyen et al., 2021; Vicidomini et al., 2021). Most studies report the transcriptome of large cell clusters of MNs, ganglion cells, neuroblasts, and glial cells due to the difficultly of matching single cell reads to a specific cell type and identity, impeding detailed analyses from scRNAseq data.

One method to deconvolve these large cell clusters is to sort cells before performing scRNAseq. For example, many labs have used the GAL4/UAS system to label and sort olfactory neurons (Li et al., 2020; McLaughlin et al., 2021; Xie et al., 2021), T cells in the fly visual system (Hörmann et al., 2020; Kurmangaliyev et al., 2020), eye-antennal discs cells (González-Blas et al., 2020) or NBs (Michki et al., 2021). On the other hand, researchers may also use the scRNAseq data to identify specific drivers, and then identify which neuron expresses this driver (Li, 2020). However, this approach reduces the scale because only a few cell types can be identified in this manner. Utilizing the expression of a gene family known to be differentially expressed within a specific subset of cells can provide a more complete examination. This expression map would generate a cell-specific atlas to annotate clusters in scRNAseq data. Here, we utilized the GAL/UAS system and showed that each MN and SN has a unique expression pattern of *dprs* and *DIPs*. Because the GAL4 is inserted in coding regions, it should capture all regulatory mechanisms and faithfully report the expression of the corresponding endogenous mRNA (Nagarkar-Jaiswal et al., 2015a, 2015b). Thus, our *dpr*/*DIP* expression could serve as a map to identify individual MNs from a MN cluster in a larval VNC sample (Nguyen et al., 2021; Vicidomini et al., 2021). In addition to *dprs* and *DIPs*, other CSP subfamilies have been reported in several scRNAseq datasets, suggesting that expression maps of other subfamilies and even combinations of subfamilies can be utilized to refine cell types in datasets (Kurmangaliyev et al., 2020; Ma et al., 2021; Xie et al., 2021).

### Insights from *dpr*/*DIP* expression maps to functional studies

The goal of developing expression maps for *dprs* and *DIPs* in MNs and SNs is to instruct the functional study of Dpr-DIP interactions. Here, we discuss potential directions based on our expression map that may serve as an entry point for future research. First, for sparsely expressed *dprs* and *DIPs*, one can use reverse genetics to analyze loss-of-function phenotypes. Most sparsely expressing *dpr/DIP-GAL4* lines are homozygous viable and hypomorphs, facilitating their use in examining phenotypes. Some *dpr/DIP-GAL4* lines are embryonic lethal, suggesting that important developmental processes are perturbed upon loss of specific *dprs* or *DIPs*. However, in these lines, we cannot rule out second site mutations so that will need to be further explored.

Another way to approach the function of Dpr-DIP interaction is focusing on the commonly or differentially expressed *dprs* and *DIPs*. For example, hierarchical clustering analyses of SNs grouped SNs from the same class together based on the expression of *dprs* and *DIPs*, suggesting that similar SNs have common *dprs* and *DIPs*. Since SNs from the same class share some downstream synaptic partners and expression of *dprs* and *DIPs*, these common CSPs may instruct recognition between SNs and interneurons. Future studies can determine the *dpr*/*DIP* expression maps in the downstream interneurons to identify synaptic partners that express cognate Dpr-DIP pairs. Instead of commonly expressed genes, differentially expressed *dprs* and *DIPs* in similar projecting neurons can shed light on connectivity mechanisms. For example, MN6-Ib and MN7-Ib identified in this study have similar morphology and innervation patterns, but with a preference for m6 and m7, respectively. One interesting question is how these neurons distinguish their muscle targets to generate such preference. Based on the expression map, MN6-Ib and MN7-Ib co-express a large subset of *dprs* and *DIPs*, but *DIP-β*, *DIP-γ*, *DIP-ε* and *dpr15* are selectively expressed. These differentially expressed genes are excellent candidates to explore the recognition mechanism of these MNs. Similar approaches can be adapted to other MNs that innervate neighboring muscles, including MN23-Ib and MN23/24-Ib or the dorsal MNs – MN9-Ib, MN10-Ib, MN1-Ib and MN2-Ib.

The Dpr-DIP interactome (Carrillo et al., 2015; Cosmanescu et al., 2018; Özkan et al., 2013) revealed promiscuity in the interactions and our expression maps showed that many cells co-express many *dprs* and *DIPs*, suggesting redundant mechanisms for synaptic recognition. Several subfamilies of CSPs are implicated in recognition, but loss-of-function mutants rarely are 100% penetrant. For example, loss of Teneurin signaling causes a 90% decrease of MN3-Ib innervation (Hong et al., 2012), and *Toll* null mutants revealed defects in 35% of MN6/7-Ib (Rose et al., 1997). These data suggested other CSPs are required in the recognition between MNs and their respective muscles. Similarly, *DIP-α* is expressed in the dorsal Is MN innervating multiple muscles, but *DIP-α* mutants only completely lose innervation of m4, suggesting that additional CSPs are required for recognition of other muscles (Ashley et al., 2019). Utilizing the *dpr*/*DIP* expression maps, co-expressed *dprs* and *DIPs* can be simultaneously knocked out to examine redundancy. Also, by combining the expression maps with scRNAseq data, additional CSPs can be identified to examine redundancy between CSP subfamilies.

### CSP expression patterns in the fly nervous system

CSPs can serve several functions in nervous system development including molecular codes for partner recognition and self-avoidance. Based on these functions, the expression of CSPs could be deterministic to instruct stereotyped synaptic connectivity or stochastic to avoid dendritic overlap and self-synapses. Thus, CSP expression patterns can suggest function. In our study, we showed that many *dprs* and *DIPs* are robustly expressed in SNs and MNs. For example, *DIP-α* and *DIP-ζ* are expressed in Is MNs across all segments (Figure 3). Several studies have implicated other CSPs in motor neuron-muscle specificity, and their expression patterns are also robust and limited to subsets of cells. For example, Capricious is expressed in MN12-Ib and some dorsal MNs (Nose, 2012; Shishido et al., 1998), and Connectin is expressed in MN27-Ib and MN29-Ib (Nose et al., 1997, 1992). Capricious and Connectin are also expressed in a unique subset of muscles. Loss-of-function and gain-of-function approaches revealed neuromuscular wiring defected, suggesting that the robust expression of Capricious and Connectin in corresponding MNs and muscles instruct synaptic partner recognition.

On the other hand, some CSPs are stochastically expressed in subsets of cells. For example, probabilistic splicing of Dscam1 generates random isoform expression in SNs to mediate dendritic self-avoidance by inhibitory homophilic interactions (Miura et al., 2013). Interestingly, we found that many *dprs* and *DIPs* are also stochastically expressed in MNs and SNs. We showed that *DIP-β* is not always expressed in dorsal da neurons (ddaC, ddaD, ddaE and ddaF) (Figure 4). Such irregular expression patterns may suggest additional functions of *dprs* and *DIPs* in circuit formation.

In this study, we also uncovered some *dprs* and *DIPs* that are expressed in a gradient along the anterior to posterior axis. Such patterns are reminiscent of the expression of several Hox genes in the VNC. For example, Ubx and Abd-A are highly expressed in anterior segments whereas Abd-B is mainly in the posterior (Estacio-Gómez and Díaz-Benjumea, 2013; Meng and Heckscher, 2020). These similar expression gradients suggest that gradient transcriptional factors may set up segment cues through *dprs* and *DIPs*.

### *dpr/DIP-GAL4* collection to enable neuron identification and manipulation

The map of *Drosophila* MNs and SNs was established decades ago using dye backfills (Broadus et al., 1995; Hoang and Chiba, 2001; Landgraf et al., 2003b). However, fluorescent dyes have some technical limitations since they do not always flow into every terminal structure, which may have resulted in some neurons being overlooked. In this study, we used a genetic approach to probe individual neurons and revealed three uncharacterized MNs – MN23-Ib, MN6-Ib (A2) and MN7-Ib (A2). Surprisingly, MN6-Ib and MN7-Ib have similar morphologies and dual innervation patterns with a preference for m6 or m7, respectively. These data suggest similar but distinct mechanisms that allow these neurons to recognize their synaptic partners.

In addition, the GAL4 lines in this study provide genetic access to manipulate subsets of neurons. In the *Drosophila* motor circuit, several studies have identified reporters that are expressed in subsets of motor neurons, muscles, and interneurons (Aponte-Santiago et al., 2020; Li et al., 2014; Pérez-Moreno and O’Kane, 2018; Wang et al., 2021). However, the coverage of these reporters is very limited (i.e. only a small number of cells can be targeted). For example, most Ib MNs cannot be individually targeted. Also, emerging evidence suggests heterogeneity of function and plasticity between different MNs (Aponte-Santiago and Littleton, 2020; Newman et al., 2017; Saunders et al., 2021), further highlighting the need for cell-specific genetic tools. To generate new genetic tools for targeting subsets of MNs, the *dpr*/*DIP* expression maps can be inspected for partially overlapping or non-overlapping *dpr/DIP-GAL4s* and converted to split-GAL4 or GAL80, respectively. For example, combining *dpr15-GAL80* and *dpr14-GAL4* should only label MN6-Ib and a few dorsal MNs; *dpr1-GAL4* and *DIP-α-GAL80* should label all Ib MNs, but not type-Is, -II ,and -III MNs; using split-GAL4, a combination of *DIP-γ-GAL4DBD* and *DIP-κ-GAL4AD* should label MN6/7-Ib and some lateral MNs. Similar approaches can be applied to SNs. Thus, the expression data in the present study and the MiMIC/CRIMIC lines provide a pipeline to expand the genetic toolbox and to label and manipulate neurons in a highly specific manner.

## Material and Methods

### Drosophila lines used in this study

All *dpr*/*DIP-GAL4* lines are listed in Table 1. Other lines used in this study are:

<u>Driver lines:</u>

*OK6-GAL4* (BL#64199)
*GMR79H07-GAL4* (gift from Troy Littleton, MIT)
*MHC-GAL80* (gift from Timothy Mosca, Thomas Jefferson University)

<u>Reporter lines:</u>

*10XUAS-mCD8::GFP* (BL#32184)
*20XUAS-mCherry* (BL# 52268)
*UAS-2XEGFP; actin-(FRT.STOP)-GAL4,UAS-FLP* (permanent reporter, gift from Ellie Heckscher, UChicago)
*UAS-nRedStinger, UAS-FLP, Ubi-p63E(FRT.STOP)-nStinger* (G-TRACE, BL#28280)
*R57C10-FLP;;UAS-MCFO* (BL#64089)

<u>Lines used to generate Trojan-*GAL4*:</u>

*yw; Sp/CyO; loxP(Trojan-GAL4)x3* (BL#60311)
*yw; loxP(Trojan-GAL4)x3; Dr/TM3,Sb,Ser* (BL#60310)
*yw,Cre,vas-phiC31:int* (BL#60299)

### Antibodies used in this study

Primary antibody:

Rabbit anti-GFP (1:40k, gift from Michael Glozter, University of Chicago)
Rabbit anti-HA (1:1000, Cell Signaling C29F4)
Mouse anti-DLG (1:100, Developmental Studies Hybridoma Bank 4F3)
Mouse anti-Repo (1:100, Developmental Studies Hybridoma Bank 8D12)
Mouse anti-Myosin (1:100, Invitrogen A31466)
Chicken anti-GFP (1:500, Invitrogen A10262)
Chicken anti-RFP (1:500, Novus Biologicals NBP2-25158)
Chicken anti-V5 (1:500, Bethyl Laboratories A190-118A)
Rat anti-Flag (1:200, Novus Biologicals NBP1-06712)

Secondary antibody:

Goat anti-Rabbit Alexa 488 (1:500, Invitrogen A11008)
Goat anti-Rabbit Alexa 568 (1:500, Invitrogen A11036)
Goat anti-Mouse Alexa 568 (1:500, Invitrogen A11031)
Goat anti-Mouse Alexa 647 (1:500, Invitrogen A32728)
Goat anti-Chicken Alexa 488 (1:500, Invitrogen A11039)
Donkey anti-Chicken Cy3 (1:500, Jackson Immunological Research 703-165-155)
Goat anti-Rat Alexa 647 (1:500, Invitrogen A21247)
Goat anti-HRP Alexa 647 (1:100, Jackson Immunological Research 123-605-021)
Goat anti-Phalloidin Alexa 405 (1:100, Invitrogen A30104)

### Fly genetics

When examining available *dpr/DIP-GAL4* lines to confirm the GAL4 insertion sites and the version of GAL4 used, we found that the original *dpr13-GAL4* no longer contained the GAL4 sequence (Barish et al., 2018; Brovero et al., 2021). Here, we generated new *dpr13-GAL4* and *dpr8-GAL4* from respective MiMIC insertion lines using Trojan exons (Diao et al., 2015). To generate *DIP-λ* CRIMIC insertions, gRNA (5’-AGCATCTATCGCTTGTGAAAGGG-3’) was designed to target the coding intron. The insertion sites and GAL4 versions are indicated in Table 1.

### qRT-PCR

Five larvae per genotype were collected and homogenized using pellet pestles (Fisher Scientific). All samples tested contained a mix of males and females, except for *dpr8-GAL4*, where only females were used due to its location on the X-chromosome and its inability to homozygous. RNA was extracted using RNAqueous Total RNA Isolation Kit (ThermoFisher AM1912) and subsequently treated with DNaseI for 30 minutes at 37°C to remove genomic DNA. cDNA was generated from 1 μg of RNA using random hexamers and SuperScript IV First-Strand Synthesis System (ThermoFisher 18091050) and remaining RNA was removed using RNase H at 37°C for 20 minutes. Primers were designed to be 18-23bp long, amplify 100-200bp, and have a melting temperature ∼60°C (Table 2). All primer locations are downstream of mapped GAL4 insertion sites and were validated with control cDNA. qRT-PCR was performed with Power SYBR Green PCR Master Mix (Bio-Rad 4368577) and run on a QuantStudio 3 (ThermoFisher). All reactions were normalized to the housekeeping gene RpL32 and control flies, yielding ΔΔCt values (Ponton et al., 2011). Relative Fold Change was calculated as 2^-ΔΔCt. Each reaction was run in technical and biological triplicates.

### Dissection and immunocytochemistry

Larval dissections and immunostaining were performed as previously described (Ashley et al., 2019). Briefly, wandering third instar larvae were dissected along the dorsal midline in PBS on a Sylgard plate and stretched out with insect pins. To visualize alary muscles, larva was dissected from the ventral side. Dissected body walls were washed once with PBS and fixed for 30min with 4% paraformaldehyde. Samples were then washed three times with PBT (PBS+0.05% TritonX100). Samples were incubated with primary antibody at 4°C overnight, washed three times with PBT, and then incubated in secondary antibody at room temperature for 2 hours. Samples were finally mounted in 30μl vectashield (Vector Laboratories). Representative images were taken with a Zeiss LSM800 confocal microscope with a 40X plan-neofluar 1.3NA objective and processed with ImageJ.

### Examining expression of *dprs* and *DIPs* in MNs and SNs

We dissected six third instar larvae from each cross and immunostained for GFP/RFP, DLG and HRP. Mounted slices were examined under Zeiss AxioImager M2 with a Lumen light engine with a 20X plan-apo 0.8NA objective. Each sample was examined twice with the same criteria to reduce human error. To map the expression of *dprs* and *DIPs* in MNs, NMJs of each MN was identified by labeling for DLG or HRP, and then examined for GFP/RFP colocalization. For expression in SNs, SN cell bodies were located by HRP, and then examined for GFP/RFP colocalization. We counted all MNs and SNs from anterior to posterior hemisegments (abdominal segment A2-A7) to gain a full *dpr*/*DIP* expression map across the body wall. Note that we did not observe the third type-Is MN (MNSNa-Is) described by (Hoang and Chiba, 2001). The pipeline and criteria of determining the expression level is below (Figure 2 – figure supplement 3):

1. In *dpr/DIP-GAL4>GFP/RFP* animals, if the reporter gene expressed constantly in a specific MN/SN in all hemisegments, then this GAL4 line is counted as “high expression level” in this MN/SN. If the fluorescent reporter is not expressed consistently in a specific MN/SN, then: (1) if the fluorescent reporter shows a gradient increase or decrease along the anterior to posterior axis, then the expression of this GAL4 line is reported as “gradient increase” or “gradient decrease”, respectively; (2) if the reporter gene does not express in a gradient, but randomly expresses in a specific MN/SN, then the expression is counted as “medium expression level” in this MN/SN. Note we did not record gradient expression for SNs, because the reporter expression had higher variation in SNs compare to MNs.
2. In the cross between *dpr/DIP-GAL4* and the permanent labeling reporter, we first confirmed the high, medium, and gradient expression level described above. Then, if a GAL4 line showed no expression in the cross to *UAS-GFP/RFP* but did show expression in the cross to the permanent labeling reporter, we counted how frequent this MN/SN is labeled: (1) if the labeling frequency is lower than 30% across all hemisegments, then this GAL4 is recorded as “low expression level” in this MN/SN because the expression could be too low to detect in the cross to *UAS-GFP/RFP* but sufficient to trigger some FLP-out; (2) if the labeling frequency is between 30%-60%, then this GAL4 expression is recorded as “medium expression level” in this MN/SN; (3) if the labeling frequency is higher than 60%, then this GAL4 expression is considered as “temporal expression” as it indicates a high GAL4 expression level temporally in early developmental stages because it triggers high frequency FLP-out. Finally, if a GAL4 is not expressed in both the cross to UAS-GFP/RFP or permanent reporter, it is recorded as “null expression”.

*dpr10-GAL4* was crossed to *UAS-GFP* together with *MHC-GAL80* to prevent muscle GFP expression, because high level of muscle GFP will mask NMJs and SN cell bodies. In addition, muscles expressing *dprs* (*dpr10* and *dpr19*) were not crossed to the permanent labeling reporter.

### Examining expression of *dprs* and *DIPs* in glial and muscles

We examined expression of *dprs* and *DIPs* in glia and muscles with the G-TRACE reporter (Evans et al., 2009). We dissected 6 larvae from each cross and immunostained for GFP, RFP, HRP, and Repo. Glial expression was confirmed by GFP/RFP colocalization with Repo. Muscle expression was confirmed by GFP/RFP positive muscle nuclei. Although the cross to *UAS-GFP/RFP* and the permanent labeling line also showed muscle expression, the diffusible GFP signal impeded the clear distinction of muscle boundaries.

### Hierarchical clustering using *dpr/DIP* expression

To perform hierarchical clustering, the expression of *dprs* and *DIPs* were first converted to binary values of “0” and “1”. Robust expression including high expression and temporal expression were considered as “1”, whereas medium and low expression, and gradient expression were considered as “0”. We reasoned that robust expression of *dprs* and *DIPs* may suggest more a significant role in the respective cell. Binary data was subjected to hierarchical analysis using Morpheus (Broad Institute) (Metric: Cosine Similarity; Method: Average). Figures were exported and color coded in Adobe Illustrator to indicate different types of MNs and SNs.

### Bouton number and dual innervation counting

To quantify m6 and m7 NMJs in wild type animals, we located Ib NMJs by DLG labeling and counted bouton number by HRP labeling. To measure the MN6-Ib or MN7-Ib NMJ sizes in *GMR79H07-GAL4>GFP* animals, we first looked for GFP colocalization with DLG to distinguish MN6-Ib and MN7-Ib. For example, if the major Ib arbor on m6 is GFP positive, then it is formed by MN6-Ib, and the GFP negative boutons are formed by MN7-Ib. We then counted the bouton numbers of each Ib arbor by HRP labeling. Statistical analyses were performed using Prism 8 software. Error bar indicates standard error of the mean (SEM).

## Acknowledgements

This work is supported by NINDS R01 NS123439 01, NSF IOS-2048080, and a UChicago Faculty Diversity Grant to R.A.C and F31NS120458 and T32 GM007183 to M.L.R. This work is also supported by funds from UChicago Biological Science Division, Committee of Developmental Biology and Department of Molecular Genetics & Cellular Biology. We thank the Drosophila Gene Disruption Project for generating MiMIC and CRIMIC insertion lines. Stocks obtained from the Bloomington Drosophila Stock Center (NIH P40OD018537) were used in this study. The hybridomas 4F3 and 8D12 were developed by Corey Goodman, and obtained from the Developmental Studies Hybridoma Bank, created by the NICHD of the NIH and maintained at The University of Iowa, Department of Biology, Iowa City, IA 52242. We would like to thank Troy Littleton (MIT), Lawrence Zipursky (UCLA) and Michelle Arbeitman (FSU) for sharing fly lines. We would also like to thank Kai Zinn, Edwin “Chip” Ferguson, Richard Fehon, Ellie Heckscher, David Pincus, Martha Plutarco Jr., and members from the Carrillo laboratory for valuable discussions and comments.

## Author contributions

Y.W. and R.A.C designed research; Y.W., M.L.R., J.A., V.A. and P.C. performed experiments; Y.W., M.L.R., and R.A.C. analyzed data; Y.W. wrote the manuscript and J.A., M.L.R., H.J.B., O.K., R.A.C. edited the manuscript.

---

**Figure 1 – figure supplement 1.**
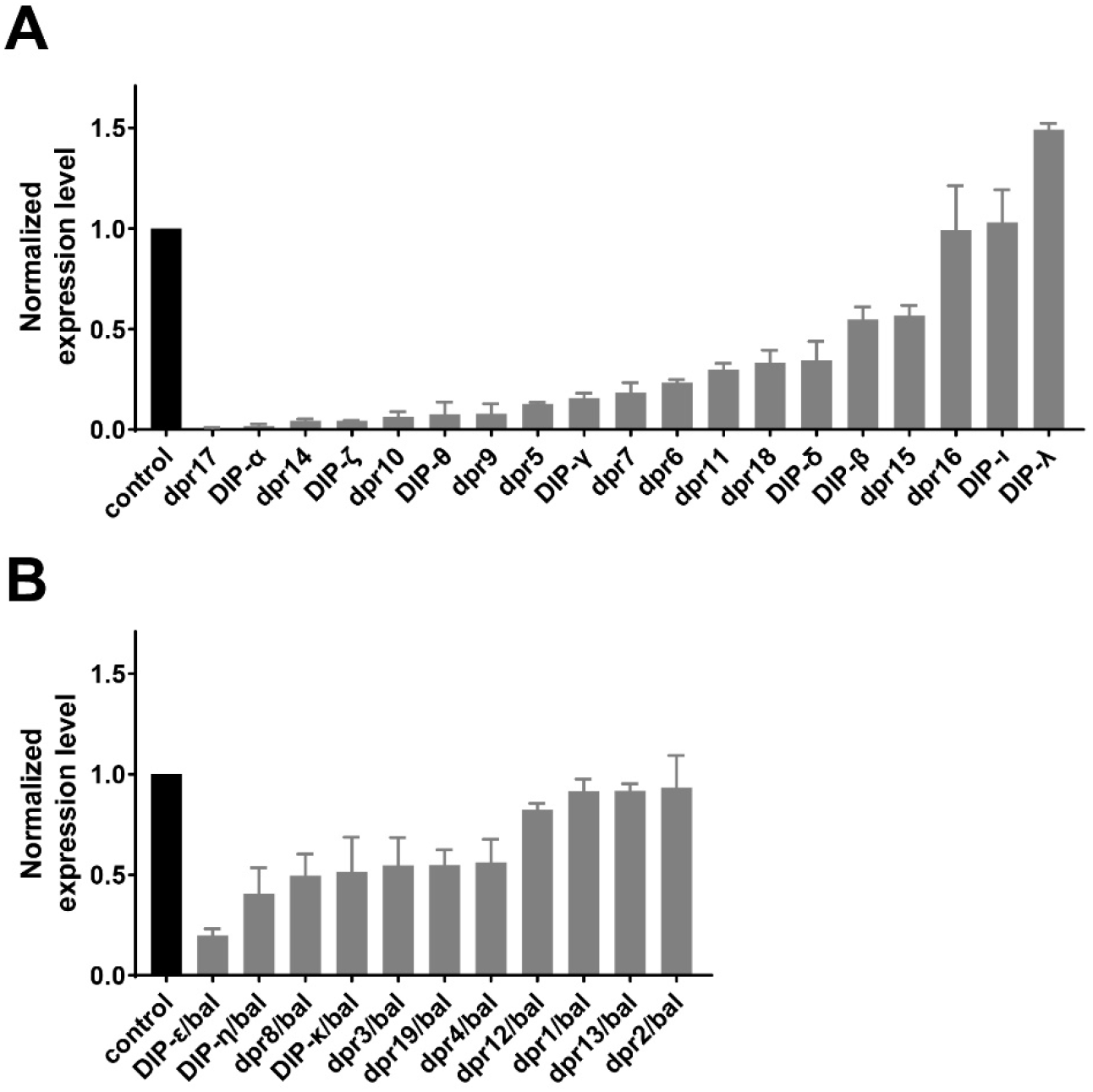
Respective mRNA level in *dpr/DIP-GAL4* lines. A. qRT-PCR results of homozygous viable *dpr/DIP-GAL4* lines. mRNA levels were double normalized to control animal and Rpl32 internal control. We found that most GAL4 lines are hypomorphs since the mRNA levels decrease below 50%. B. qRT-PCR results of homozygous lethal *dpr/DIP-GAL4* lines. mRNA levels were double normalized to control animal and Rpl32 internal control. We found that most GAL4 lines have an expression level near 50%, indicating that they are hypomorphs.

**Figure 2 – figure supplement 1.**
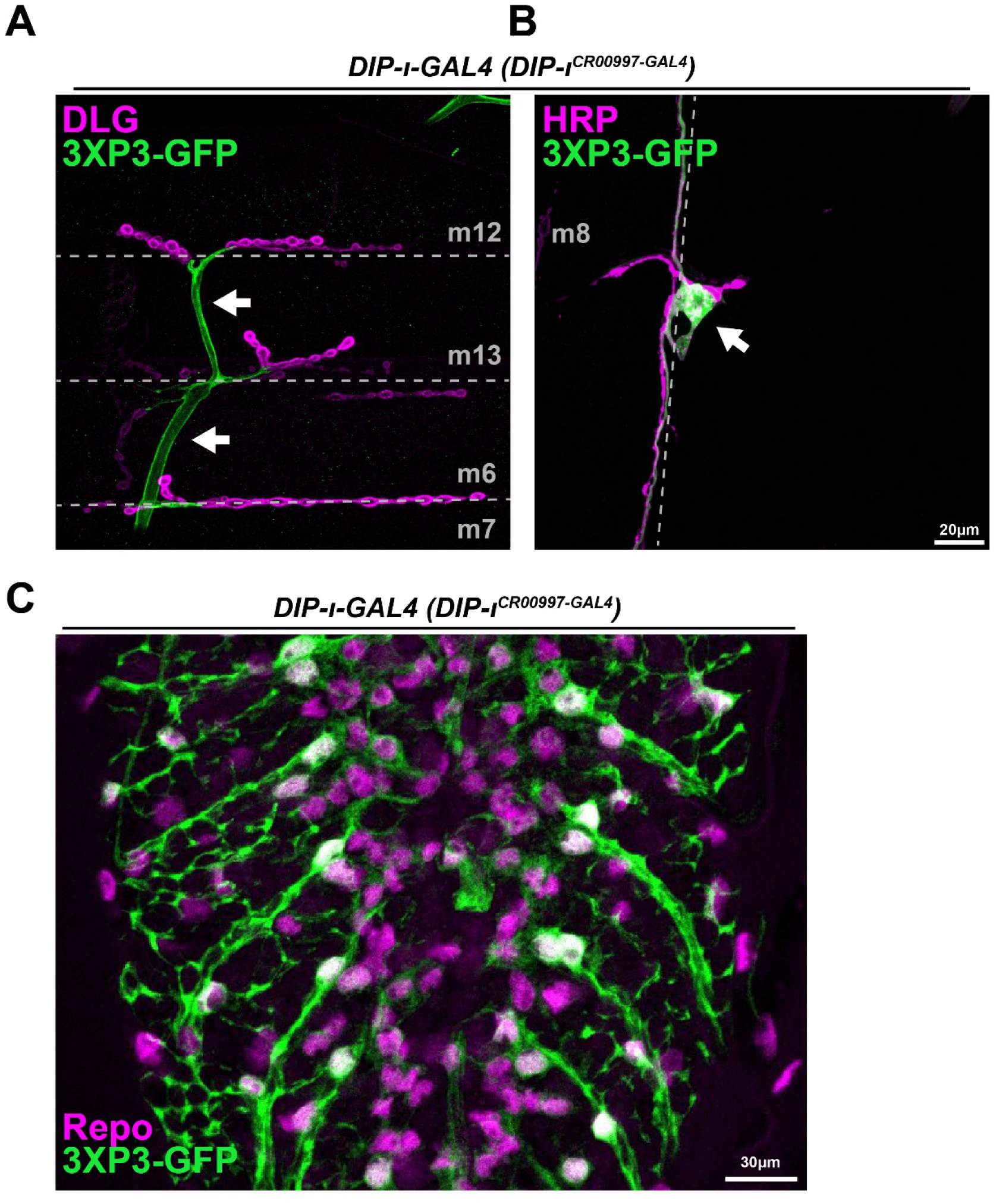
Expression of CRIMIC 3XP3-GFP in nervous system. A-B. Representative images labeled with DLG (magenta) and GFP (green) showing the expression of 3XP3-GFP in peripheral glial cells (arrow) and the lbd SN (arrow). C. Representative image labeled with the glial cell marker, Repo (magenta), and GFP (green). Note the expression of 3XP3-GFP in some glial cells in the VNC.

**Figure 2 – figure supplement 2.**
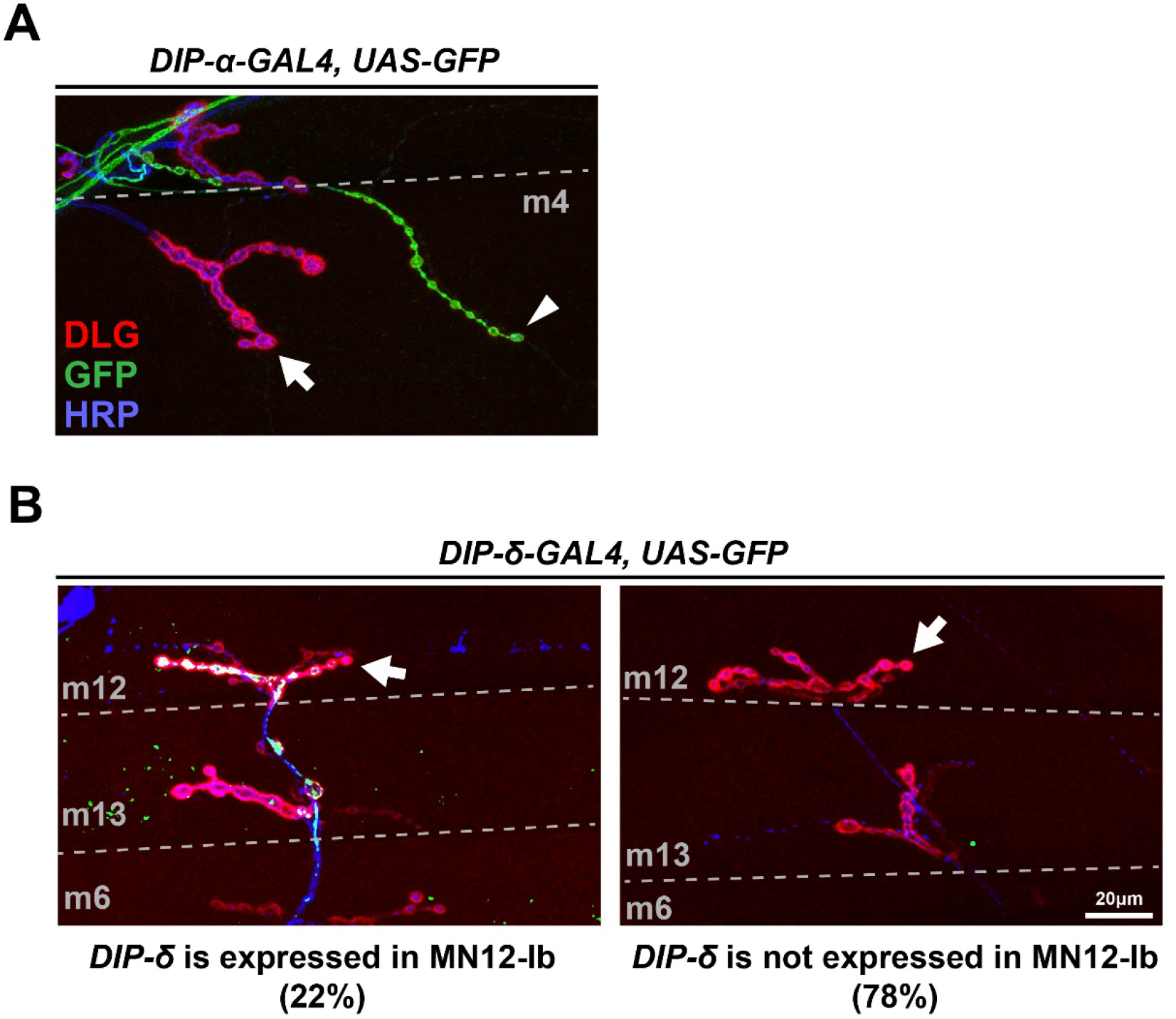
Expression of *DIP-α* and *DIP-δ* in MNs. A. Representative image labeled for HRP (blue), DLG (red) and GFP (green) showing the expression of *DIP-α* in Is MNJ (arrowhead) but not in adjacent Ib NMJ (arrow). B. Representative images labeled with HRP (blue), DLG (red) and GFP (green) showing the varied expression of *DIP-δ* in MN12-Ib (arrow). Note that 22% MN12-Ib express *DIP-δ* (left) whereas 78% do not express (right).

**Figure 2 – figure supplement 3.**
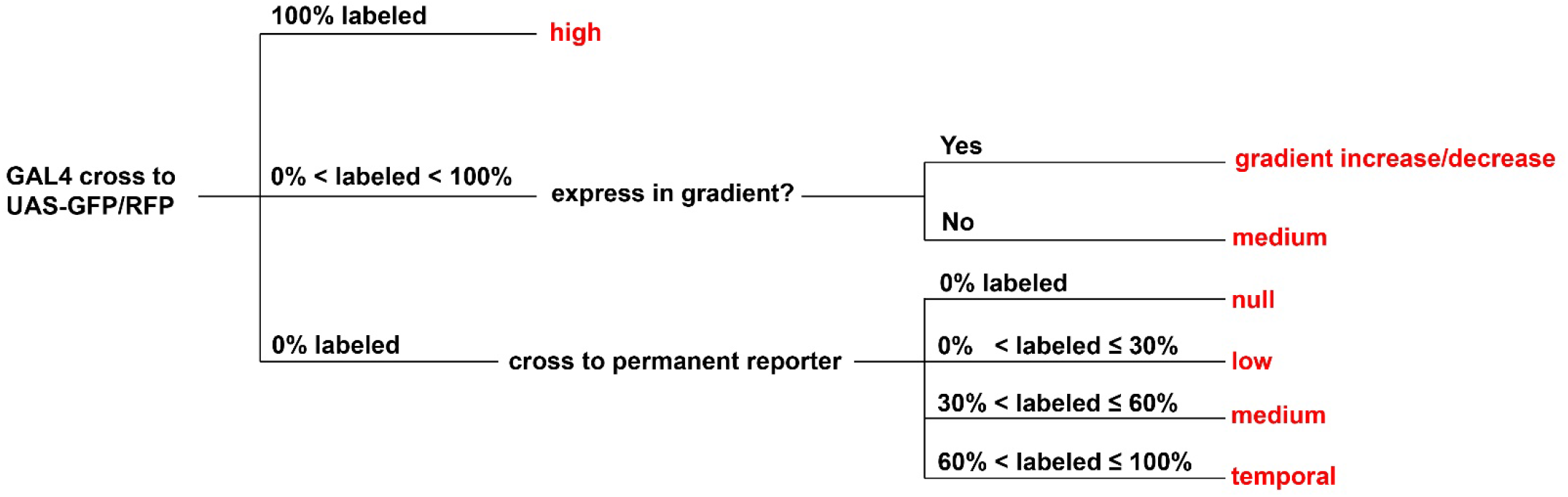
Criteria to determine the expression of GAL4 in a certain MN/SN. A graphical flow chart depicting how we scored the *dpr*/*DIP* expression data into categories.

**Figure 3 – figure supplement 1.**
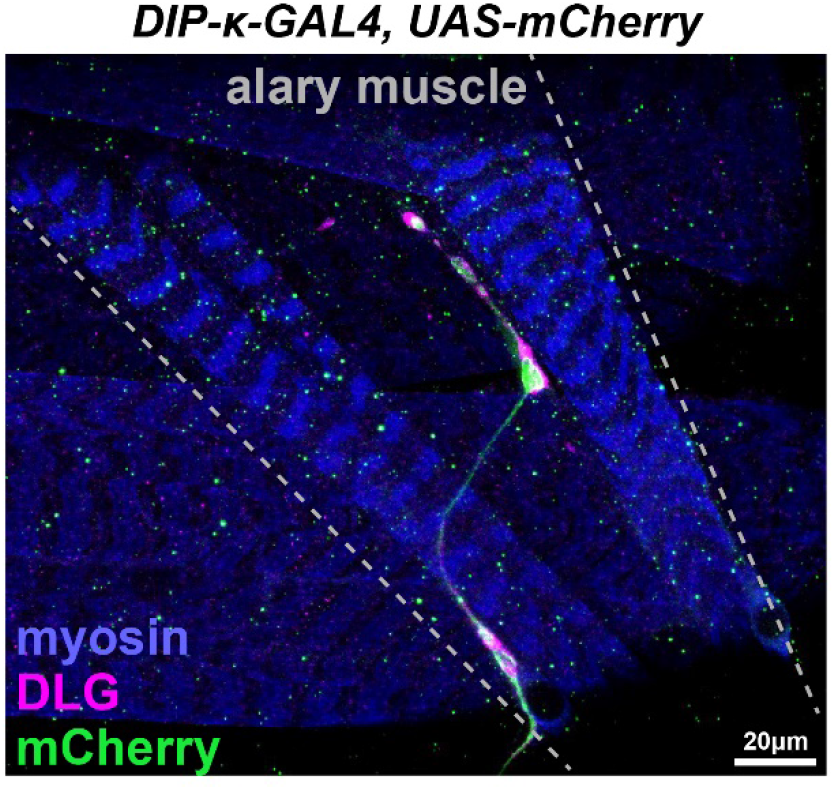
The alary MN also expresses *dprs* and *DIPs*. Representative image labeled for myosin (blue), DLG (magenta) and mCherry (green) showing *DIP-κ* expression in the alary MN. Alary MNs display features of excitatory MNs as they have DLG accumulation around boutons.

**Figure 4 – figure supplement 1.**
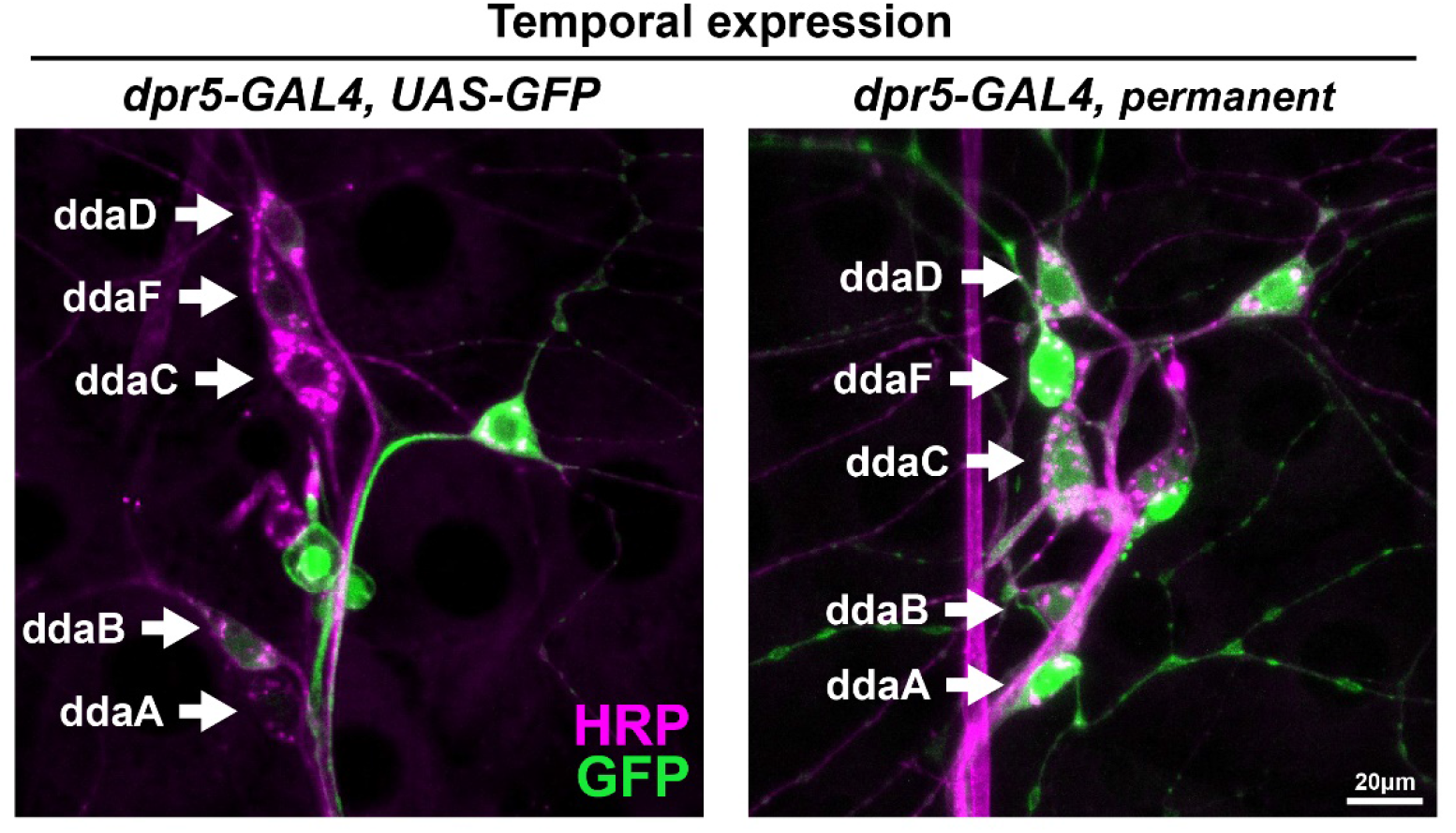
Temporal expression of *dpr5* in the dorsal da neuron cluster. ddaA, ddaC, ddaF and ddaD are not labeled in *dpr5-GAL4>GFP* larvae but are robustly labeled in the cross to the permanent reporter. Therefore, *dpr5* is temporally expressed in these SNs. ddaB is labeled in the cross to the real-time reporter with a low frequency, thus *dpr5* is considered as low expression in ddaB.

**Figure 7 – figure supplement 1.**
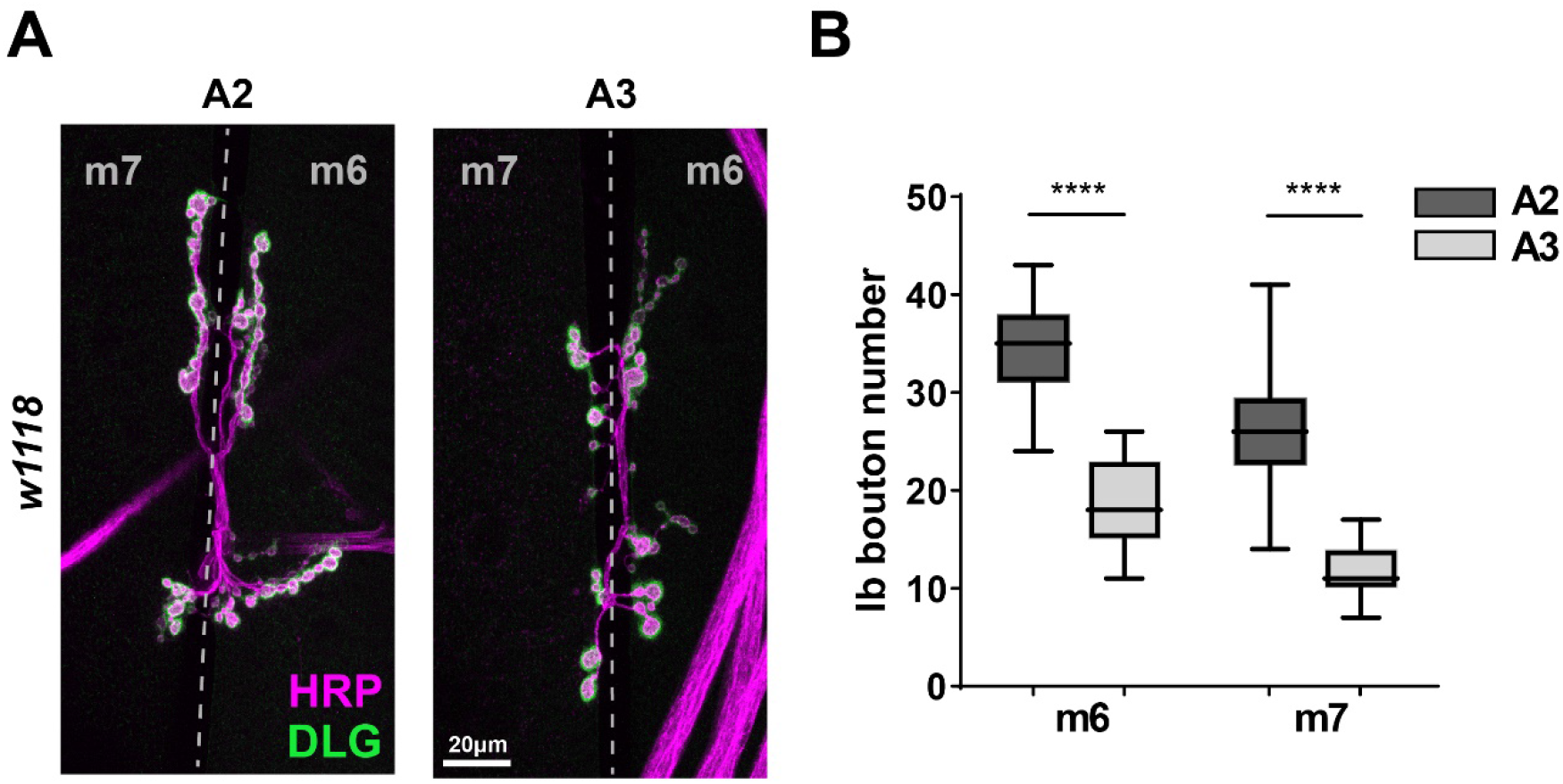
Larger NMJs on m6 and m7 in A2. A. Representative images showing larger type-Ib NMJs on m6 and m7 in A2 compared to A3. B. Bouton number counts from m6 and m7 in A2 and A3 confirmed that A2 NMJs are double the size of A3.

**Figure 8 – figure supplement 1.**
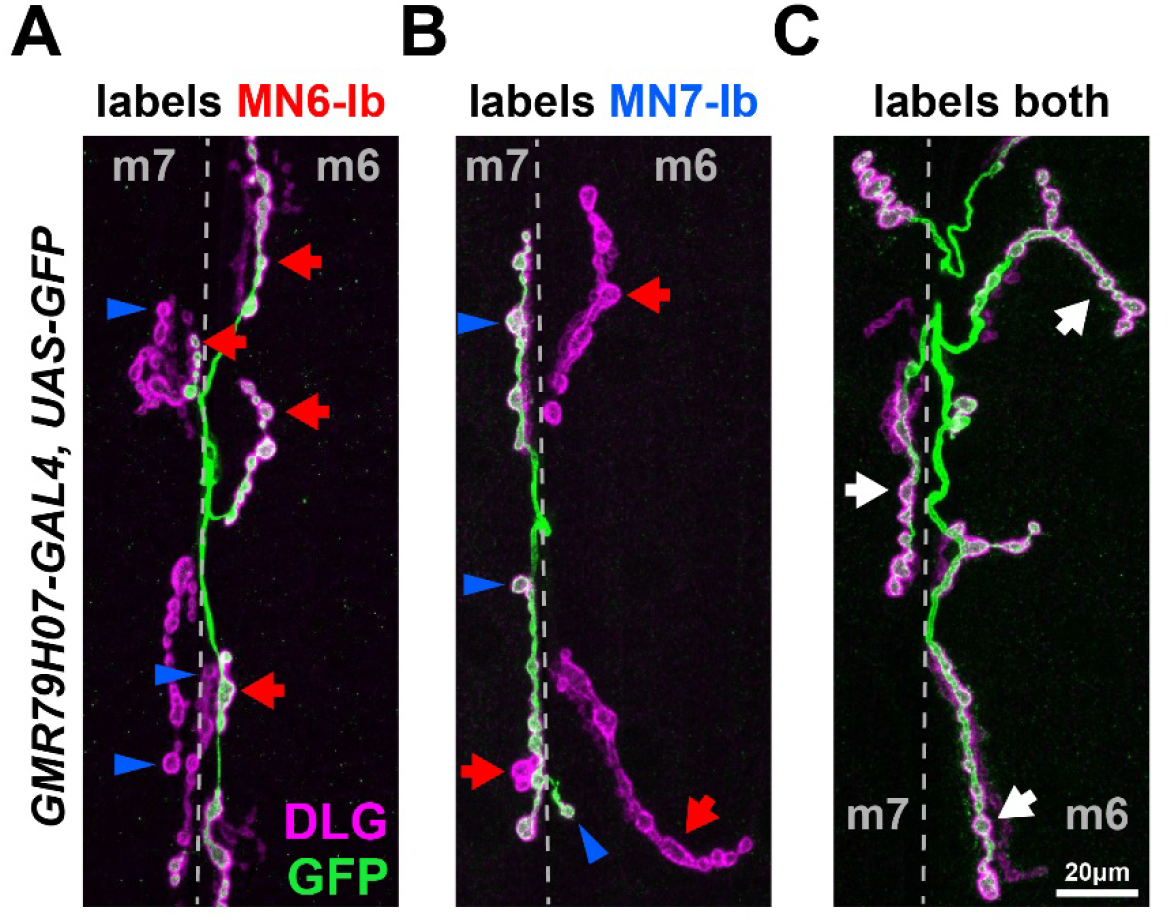
*GMR79H07-GAL4* randomly labels MN6-Ib and MN7-Ib. A previous study reported that *GMR79H07-GAL4* labels type-Ib NMJs on m6 in A2 (Aponte-Santiago et al., 2020). We crossed this driver to *UAS-GFP* and found inconsistent expression patterns since it (A) sometimes only labels MN6-Ib (red arrows), (B) sometimes only labels MN7-Ib (blue arrowheads), or (C) sometimes labels both MNs (white arrows). These expression patterns support the existence of MN6-Ib and MN7-Ib and their dual innervation properties.

**Figure 9 – figure supplement 1.**
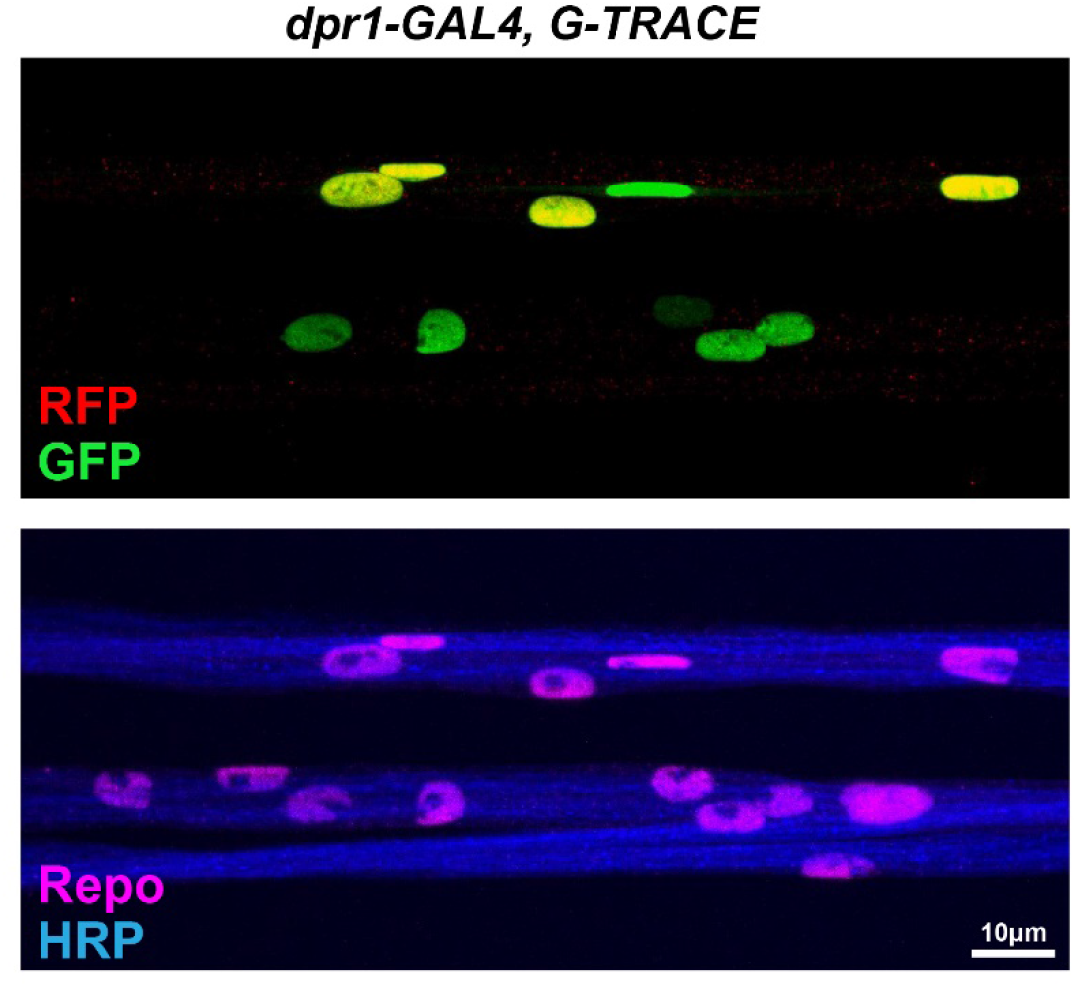
The G-TRACE system revealed *dpr1* expression in peripheral glial cells. *dpr1* is the only *dpr/DIP* expressed in peripheral glial cells. *dpr1-GAL4* is expressed in subsets of peripheral glial cells as indicated by some glial nuclei labeled by both GFP and RFP, some only by GFP, and some lacking both GFP and RFP.

**Figure 9 – figure supplement 2.**
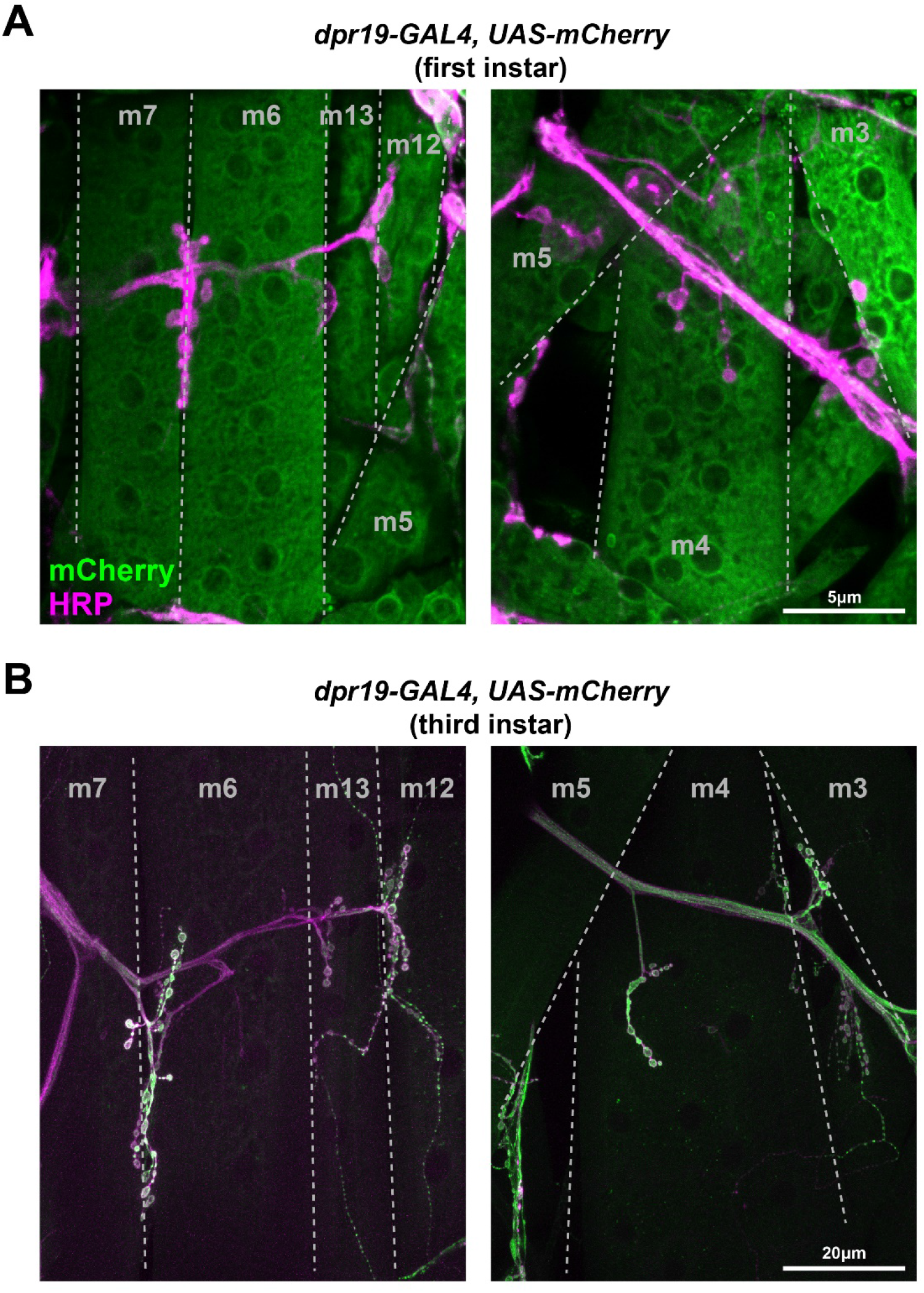
*dpr19* is expressed in muscles in early larval development. A. Representative images of *dpr19-GAL4>mCherry* in first instar larvae. *dpr19* is expressed in ventral (left) and dorsal (right) muscles. B. Representative images of *dpr19-GAL4>mCherry* in third instar larvae. *dpr19* is not expressed in ventral (left) or dorsal (right) muscles.

## References

Allen AM, Neville MC, Birtles S, Croset V, Treiber CD, Waddell S, Goodwin SF. 2020. A single-cell transcriptomic atlas of the adult Drosophila ventral nerve cord. Elife 9:e54074. doi:10.7554/elife.54074

Aponte-Santiago NA, Littleton JT. 2020. Synaptic Properties and Plasticity Mechanisms of Invertebrate Tonic and Phasic Neurons. Frontiers in Physiology 11:611982. doi:10.3389/fphys.2020.611982

Aponte-Santiago NA, Ormerod KG, Akbergenova Y, Littleton JT. 2020. Synaptic plasticity induced by differential manipulation of tonic and phasic motoneurons in Drosophila. J Neurosci Official J Soc Neurosci JN-RM-0925-20. doi:10.1523/jneurosci.0925-20.2020

Ariss MM, Islam ABMMK, Critcher M, Zappia MP, Frolov MV. 2018. Single cell RNA-sequencing identifies a metabolic aspect of apoptosis in Rbf mutant. Nat Commun 9:5024. doi:10.1038/s41467-018-07540-z

Ashley J, Sorrentino V, Lobb-Rabe M, Nagarkar-Jaiswal S, Tan L, Xu S, Xiao Q, Zinn K, Carrillo RA. 2019. Transsynaptic interactions between IgSF proteins DIP-α and Dpr10 are required for motor neuron targeting specificity. eLife 8:e42690. doi:10.7554/eLife.42690

Avalos CB, Maier GL, Bruggmann R, Sprecher SG. 2019. Single cell transcriptome atlas of the Drosophila larval brain. Elife 8:e50354. doi:10.7554/elife.50354

Barish S, Nuss S, Strunilin I, Bao S, Mukherjee S, Jones CD, Volkan PC. 2018. Combinations of DIPs and Dprs control organization of olfactory receptor neuron terminals in Drosophila. Plos Genet 14:e1007560. doi:10.1371/journal.pgen.1007560

Bataillé L, Frendo J-L, Vincent A. 2015. Hox control of Drosophila larval anatomy; The Alary and Thoracic Alary-Related Muscles. Mech Develop 138:170–176. doi:10.1016/j.mod.2015.07.005

Bate M. 1990. The embryonic development of larval muscles in Drosophila. Dev Camb Engl 110:791–804.

Bittern J, Pogodalla N, Ohm H, Brüser L, Kottmeier R, Schirmeier S, Klämbt C. 2020. Neuron-Glia Interaction in the Drosophila nervous system. Dev Neurobiol. doi:10.1002/dneu.22737

Bornstein B, Meltzer H, Adler R, Alyagor I, Berkun V, Cummings G, Reh F, Keren-Shaul H, David E, Riemensperger T, Schuldiner O. 2021. Transneuronal Dpr12/DIP-δ interactions facilitate compartmentalized dopaminergic innervation of Drosophila mushroom body axons. Embo J e105763. doi:10.15252/embj.2020105763

Boukhatmi H, Schaub C, Bataillé L, Reim I, Frendo J-L, Frasch M, Vincent A. 2014. An Org-1–Tup transcriptional cascade reveals different types of alary muscles connecting internal organs in Drosophila. Development 141:3761–3771. doi:10.1242/dev.111005

Broadus J, Skeath JB, Spana EP, Bossing T, Technau G, Doe CQ. 1995. New neuroblast markers and the origin of the aCC/pCC neurons in the Drosophila central nervous system. Mech Develop 53:393–402. doi:10.1016/0925-4773(95)00454-8

Brovero SG, Fortier JC, Hu H, Lovejoy PC, Newell NR, Palmateer CM, Tzeng R-Y, Lee P-T, Zinn K, Arbeitman MN. 2021. Investigation of Drosophila fruitless neurons that express Dpr/DIP cell adhesion molecules. Elife 10:e63101. doi:10.7554/elife.63101

Carrillo RA, Özkan E, Menon KP, Nagarkar-Jaiswal S, Lee P-T, Jeon M, Birnbaum ME, Bellen HJ, Garcia KC, Zinn K. 2015. Control of Synaptic Connectivity by a Network of Drosophila IgSF Cell Surface Proteins. Cell 163:1770–1782. doi:10.1016/j.cell.2015.11.022

Cheng S, Park Y, Kurleto JD, Jeon M, Zinn K, Thornton JW, Özkan E. 2019. Family of neural wiring receptors in bilaterians defined by phylogenetic, biochemical, and structural evidence. Proc National Acad Sci 116:201818631. doi:10.1073/pnas.1818631116

Chiba A, Snow P, Keshishian H, Hotta Y. 1995. Fasciclin III as a synaptic target recognition molecule in Drosophila. Nature 374:166–168. doi:10.1038/374166a0

Choi JC, Park D, Griffith LC. 2004. Electrophysiological and Morphological Characterization of Identified Motor Neurons in the Drosophila Third Instar Larva Central Nervous System. J Neurophysiol 91:2353–2365. doi:10.1152/jn.01115.2003

Cosmanescu F, Katsamba PS, Sergeeva AP, Ahlsen G, Patel SD, Brewer JJ, Tan L, Xu S, Xiao Q, Nagarkar-Jaiswal S, Nern A, Bellen HJ, Zipursky SL, Honig B, Shapiro L. 2018. Neuron-Subtype-Specific Expression, Interaction Affinities, and Specificity Determinants of DIP/Dpr Cell Recognition Proteins. Neuron 100:1385–1400.e6. doi:10.1016/j.neuron.2018.10.046

Courgeon M, Desplan C. 2019. Coordination between stochastic and deterministic specification in the Drosophila visual system. Science 366:eaay6727. doi:10.1126/science.aay6727

Davie K, Janssens J, Koldere D, Waegeneer MD, Pech U, Kreft Ł, Aibar S, Makhzami S, Christiaens V, González-Blas CB, Poovathingal S, Hulselmans G, Spanier KI, Moerman T, Vanspauwen B, Geurs S, Voet T, Lammertyn J, Thienpont B, Liu S, Konstantinides N, Fiers M, Verstreken P, Aerts S. 2018. A Single-Cell Transcriptome Atlas of the Aging Drosophila Brain. Cell 174:982–998.e20. doi:10.1016/j.cell.2018.05.057

Davis GW, Schuster CM, Goodman CS. 1997. Genetic Analysis of the Mechanisms Controlling Target Selection: Target-Derived Fasciclin II Regulates the Pattern of Synapse Formation. Neuron 19:561–573. doi:10.1016/s0896-6273(00)80372-4

Diao Fengqiu, Ironfield H, Luan H, Diao Feici, Shropshire WC, Ewer J, Marr E, Potter CJ, Landgraf M, White BH. 2015. Plug-and-Play Genetic Access to Drosophila Cell Types using Exchangeable Exon Cassettes. Cell Reports 10:1410–1421. doi:10.1016/j.celrep.2015.01.059

Doe CQ, Smouse D, Goodman CS. 1988. Control of neuronal fate by the Drosophila segmentation gene even-skipped. Nature 333:376–378. doi:10.1038/333376a0

Duan X, Krishnaswamy A, De la Huerta I, Sanes JR. 2014. Type II Cadherins Guide Assembly of a Direction-Selective Retinal Circuit. Cell 158:793–807. doi:10.1016/j.cell.2014.06.047

Duan X, Krishnaswamy A, Laboulaye MA, Liu J, Peng Y-R, Yamagata M, Toma K, Sanes JR. 2018. Cadherin Combinations Recruit Dendrites of Distinct Retinal Neurons to a Shared Interneuronal Scaffold. Neuron 99:1145–1154.e6. doi:10.1016/j.neuron.2018.08.019

Estacio-Gómez A, Díaz-Benjumea FJ. 2013. Roles of Hox genes in the patterning of the central nervous system of Drosophila. Fly 8:26–32. doi:10.4161/fly.27424

Evans CJ, Olson JM, Ngo KT, Kim E, Lee NE, Kuoy E, Patananan AN, Sitz D, Tran P, Do M-T, Yackle K, Cespedes A, Hartenstein V, Call GB, Banerjee U. 2009. G-TRACE: rapid Gal4-based cell lineage analysis in Drosophila. Nat Methods 6:603–605. doi:10.1038/nmeth.1356

Garrett AM, Khalil A, Walton DO, Burgess RW. 2018. DSCAM promotes self-avoidance in the developing mouse retina by masking the functions of cadherin superfamily members. Proc National Acad Sci 115:201809430. doi:10.1073/pnas.1809430115

Genovese S, Clément R, Gaultier C, Besse F, Narbonne-Reveau K, Daian F, Foppolo S, Luis NM, Maurange C. 2019. Coopted temporal patterning governs cellular hierarchy, heterogeneity and metabolism in Drosophila neuroblast tumors. Elife 8:e50375. doi:10.7554/elife.50375

González-Blas CB, Quan X, Duran-Romaña R, Taskiran II, Koldere D, Davie K, Christiaens V, Makhzami S, Hulselmans G, Waegeneer M, Mauduit D, Poovathingal S, Aibar S, Aerts S. 2020. Identification of genomic enhancers through spatial integration of single-cell transcriptomics and epigenomics. Mol Syst Biol 16:e9438. doi:10.15252/msb.20209438

Gorczyca MG, Phillis RW, Budnik V. 1994. The role of tinman, a mesodermal cell fate gene, in axon pathfinding during the development of the transverse nerve in Drosophila. Dev Camb Engl 120:2143–52.

Grueber WB, Jan LY, Jan YN. 2002. Tiling of the Drosophila epidermis by multidendritic sensory neurons. Dev Camb Engl 129:2867–78.

Grueber WB, Ye B, Yang C-H, Younger S, Borden K, Jan LY, Jan Y-N. 2007. Projections of Drosophila multidendritic neurons in the central nervous system: links with peripheral dendrite morphology. Development 134:55–64. doi:10.1242/dev.02666

Guan B, Hartmann B, Kho Y-H, Gorczyca M, Budnik V. 1996. The Drosophila tumor suppressor gene, dlg, is involved in structural plasticity at a glutamatergic synapse. Curr Biol 6:695–706. doi:10.1016/s0960-9822(09)00451-5

Hattori D, Chen Y, Matthews BJ, Salwinski L, Sabatti C, Grueber WB, Zipursky SL. 2009. Robust discrimination between self and non-self neurites requires thousands of Dscam1 isoforms. Nature 461:644–648. doi:10.1038/nature08431

Heckscher ES, Zarin AA, Faumont S, Clark MQ, Manning L, Fushiki A, Schneider-Mizell CM, Fetter RD, Truman JW, Zwart MF, Landgraf M, Cardona A, Lockery SR, Doe CQ. 2015. Even-Skipped+ Interneurons Are Core Components of a Sensorimotor Circuit that Maintains Left-Right Symmetric Muscle Contraction Amplitude. Neuron 88:314–329. doi:10.1016/j.neuron.2015.09.009

Hoang B, Chiba A. 2001. Single-Cell Analysis of Drosophila Larval Neuromuscular Synapses. Dev Biol 229:55–70. doi:10.1006/dbio.2000.9983

Hong W, Mosca TJ, Luo L. 2012. Teneurins instruct synaptic partner matching in an olfactory map. Nature 484:201–207. doi:10.1038/nature10926

Honig B, Shapiro L. 2020. Adhesion Protein Structure, Molecular Affinities, and Principles of Cell-Cell Recognition. Cell 181:520–535. doi:10.1016/j.cell.2020.04.010

Hooper JE. 1986. Homeotic gene function in the muscles of Drosophila larvae. Embo J 5:2321–2329. doi:10.1002/j.1460-2075.1986.tb04500.x

Hörmann N, Schilling T, Ali AH, Serbe E, Mayer C, Borst A, Pujol-Martí J. 2020. A combinatorial code of transcription factors specifies subtypes of visual motion-sensing neurons in Drosophila. Development 147:dev186296. doi:10.1242/dev.186296

Inaki M, Shinza-Kameda M, Ismat A, Frasch M, Nose A. 2010. Drosophila Tey represses transcription of the repulsive cue Toll and generates neuromuscular target specificity. Dev Camb Engl 137:2139–46. doi:10.1242/dev.046672

Jan LY, Jan YN. 1982. Antibodies to horseradish peroxidase as specific neuronal markers in Drosophila and in grasshopper embryos. Proc National Acad Sci 79:2700–2704. doi:10.1073/pnas.79.8.2700

Jontes JD. 2017. The Cadherin Superfamily in Neural Circuit Assembly. Csh Perspect Biol 10:a029306. doi:10.1101/cshperspect.a029306

Kanca O, Zirin J, Garcia-Marques J, Knight SM, Yang-Zhou D, Amador G, Chung H, Zuo Z, Ma L, He Y, Lin W-W, Fang Y, Ge M, Yamamoto S, Schulze KL, Hu Y, Spradling AC, Mohr SE, Perrimon N, Bellen HJ. 2019. An efficient CRISPR-based strategy to insert small and large fragments of DNA using short homology arms. Elife 8:e51539. doi:10.7554/elife.51539

Kim MD, Wen Y, Jan Y-N. 2009. Patterning and organization of motor neuron dendrites in the Drosophila larva. Dev Biol 336:213–221. doi:10.1016/j.ydbio.2009.09.041

Konstantinides N, Kapuralin K, Fadil C, Barboza L, Satija R, Desplan C. 2018. Phenotypic Convergence: Distinct Transcription Factors Regulate Common Terminal Features. Cell 174:622–635.e13. doi:10.1016/j.cell.2018.05.021

Kose H, Rose D, Zhu X, Chiba A. 1997. Homophilic synaptic target recognition mediated by immunoglobulin-like cell adhesion molecule Fasciclin III. Dev Camb Engl 124:4143–52.

Kottmeier R, Bittern J, Schoofs A, Scheiwe F, Matzat T, Pankratz M, Klämbt C. 2020. Wrapping glia regulates neuronal signaling speed and precision in the peripheral nervous system of Drosophila. Nat Commun 11:4491. doi:10.1038/s41467-020-18291-1

Krishnaswamy A, Yamagata M, Duan X, Hong YK, Sanes JR. 2015. Sidekick 2 directs formation of a retinal circuit that detects differential motion. Nature 524:466–470. doi:10.1038/nature14682

Kurmangaliyev YZ, Yoo J, Valdes-Aleman J, Sanfilippo P, Zipursky SL. 2020. Transcriptional Programs of Circuit Assembly in the Drosophila Visual System. Neuron 108. doi:10.1016/j.neuron.2020.10.006

Kurusu M, Cording A, Taniguchi M, Menon K, Suzuki E, Zinn K. 2008. A screen of cell-surface molecules identifies leucine-rich repeat proteins as key mediators of synaptic target selection. Neuron 59:972–85. doi:10.1016/j.neuron.2008.07.037

Kwon JY, Dahanukar A, Weiss LA, Carlson JR. 2014. A map of taste neuron projections in the Drosophila CNS. J Biosciences 39:565–574. doi:10.1007/s12038-014-9448-6

Landgraf M, Bossing T, Technau GM, Bate M. 1997. The Origin, Location, and Projections of the Embryonic Abdominal Motorneurons of Drosophila. J Neurosci 17:9642–9655. doi:10.1523/jneurosci.17-24-09642.1997

Landgraf M, Jeffrey V, Fujioka M, Jaynes JB, Bate M. 2003a. Embryonic Origins of a Motor System: Motor Dendrites Form a Myotopic Map in Drosophila. Plos Biol 1:e41. doi:10.1371/journal.pbio.0000041

Landgraf M, Sánchez-Soriano N, Technau GM, Urban J, Prokop A. 2003b. Charting the Drosophila neuropile: a strategy for the standardised characterisation of genetically amenable neurites. Dev Biol 260:207–225. doi:10.1016/s0012-1606(03)00215-x

Landgraf M, Thor S. 2006. Development of Drosophila motoneurons: Specification and morphology. Semin Cell Dev Biol 17:3–11. doi:10.1016/j.semcdb.2005.11.007

Lee P-T, Zirin J, Kanca O, Lin W-W, Schulze KL, Li-Kroeger D, Tao R, Devereaux C, Hu Y, Chung V, Fang Y, He Y, Pan H, Ge M, Zuo Z, Housden BE, Mohr SE, Yamamoto S, Levis RW, Spradling AC, Perrimon N, Bellen HJ. 2018. A gene-specific T2A-GAL4 library for Drosophila. Elife 7:e35574. doi:10.7554/elife.35574

Li H. 2020. Single-cell RNA sequencing in Drosophila: Technologies and applications. Wiley Interdiscip Rev Dev Biology e396. doi:10.1002/wdev.396

Li H, Li T, Horns F, Li J, Xie Q, Xu C, Wu B, Kebschull JM, McLaughlin CN, Kolluru SS, Jones RC, Vacek D, Xie A, Luginbuhl DJ, Quake SR, Luo L. 2020. Single-Cell Transcriptomes Reveal Diverse Regulatory Strategies for Olfactory Receptor Expression and Axon Targeting. Curr Biol 30:1189–1198.e5. doi:10.1016/j.cub.2020.01.049

Li H-H, Kroll JR, Lennox SM, Ogundeyi O, Jeter J, Depasquale G, Truman JW. 2014. A GAL4 Driver Resource for Developmental and Behavioral Studies on the Larval CNS of Drosophila. Cell Reports 8:897–908. doi:10.1016/j.celrep.2014.06.065

Lnenicka GA, Keshishian H. 2000. Identified motor terminals in Drosophila larvae show distinct differences in morphology and physiology. J Neurobiol 43:186–197. doi:10.1002/(sici)1097-4695(200005)43:2<186::aid-neu8>3.0.co;2-n

Logan J, Falck-Pedersen E, Darnell JE, Shenk T. 1987. A poly(A) addition site and a downstream termination region are required for efficient cessation of transcription by RNA polymerase II in the mouse beta maj-globin gene. Proc National Acad Sci 84:8306–8310. doi:10.1073/pnas.84.23.8306

Ma D, Przybylski D, Abruzzi KC, Schlichting M, Li Q, Long X, Rosbash M. 2021. A transcriptomic taxonomy of Drosophila circadian neurons around the clock. Elife 10:e63056. doi:10.7554/elife.63056

Macleod GT, Suster ML, Charlton MP, Atwood HL. 2003. Single neuron activity in the Drosophila larval CNS detected with calcium indicators. J Neurosci Meth 127:167–178. doi:10.1016/s0165-0270(03)00127-4

Mauss A, Tripodi M, Evers JF, Landgraf M. 2009. Midline Signalling Systems Direct the Formation of a Neural Map by Dendritic Targeting in the Drosophila Motor System. Plos Biol 7:e1000200. doi:10.1371/journal.pbio.1000200

McLaughlin CN, Brbić M, Xie Q, Li T, Horns F, Kolluru SS, Kebschull JM, Vacek D, Xie A, Li J, Jones RC, Leskovec J, Quake SR, Luo L, Li H. 2021. Single-cell transcriptomes of developing and adult olfactory receptor neurons in Drosophila. Elife 10:e63856. doi:10.7554/elife.63856

Meltzer S, Yadav S, Lee J, Soba P, Younger SH, Jin P, Zhang W, Parrish J, Jan LY, Jan Y-N. 2016. Epidermis-Derived Semaphorin Promotes Dendrite Self-Avoidance by Regulating Dendrite-Substrate Adhesion in Drosophila Sensory Neurons. Neuron 89:741–755. doi:10.1016/j.neuron.2016.01.020

Meng JL, Heckscher ES. 2020. Development of motor circuits: From neuronal stem cells and neuronal diversity to motor circuit assembly. Curr Top Dev Biol. doi:10.1016/bs.ctdb.2020.11.010

Menon KP, Carrillo RA, Zinn K. 2013. Development and plasticity of the Drosophila larval neuromuscular junction. Wiley Interdiscip Rev Dev Biology 2:647–70. doi:10.1002/wdev.108

Menon KP, Kulkarni V, Takemura S, Anaya M, Zinn K. 2019. Interactions between Dpr11 and DIP-γ control selection of amacrine neurons in Drosophila color vision circuits. Elife 8:e48935. doi:10.7554/elife.48935

Merritt D, Whitington P. 1995. Central projections of sensory neurons in the Drosophila embryo correlate with sensory modality, soma position, and proneural gene function. J Neurosci 15:1755–1767. doi:10.1523/jneurosci.15-03-01755.1995

Merritt DJ, Murphey RK. 1992. Projections of leg proprioceptors within the CNS of the fly Phormia in relation to the generalized insect ganglion. J Comp Neurol 322:16–34. doi:10.1002/cne.903220103

Michki NS, Li Y, Sanjasaz K, Zhao Y, Shen FY, Walker LA, Cao W, Lee C-Y, Cai D. 2021. The molecular landscape of neural differentiation in the developing Drosophila brain revealed by targeted scRNA-seq and multi-informatic analysis. Cell Reports 35:109039. doi:10.1016/j.celrep.2021.109039

Miura SK, Martins A, Zhang KX, Graveley BR, Zipursky SL. 2013. Probabilistic Splicing of Dscam1 Establishes Identity at the Level of Single Neurons. Cell 155:1166–1177. doi:10.1016/j.cell.2013.10.018

Murphey RK, Possidente D, Pollack G, Merritt DJ. 1989. Modality-specific axonal projections in the CNS of the flies Phormia and Drosophila. J Comp Neurol 290:185–200. doi:10.1002/cne.902900203

Nagarkar-Jaiswal S, DeLuca SZ, Lee P-T, Lin W-W, Pan H, Zuo Z, Lv J, Spradling AC, Bellen HJ. 2015a. A genetic toolkit for tagging intronic MiMIC containing genes. Elife 4:e08469. doi:10.7554/elife.08469

Nagarkar-Jaiswal S, Lee P-T, Campbell ME, Chen K, Anguiano-Zarate S, Gutierrez MC, Busby T, Lin W-W, He Y, Schulze KL, Booth BW, Evans-Holm M, Venken KJ, Levis RW, Spradling AC, Hoskins RA, Bellen HJ. 2015b. A library of MiMICs allows tagging of genes and reversible, spatial and temporal knockdown of proteins in Drosophila. Elife 4:e05338. doi:10.7554/elife.05338

Nern A, Pfeiffer BD, Rubin GM. 2015. Optimized tools for multicolor stochastic labeling reveal diverse stereotyped cell arrangements in the fly visual system. Proc National Acad Sci 112:E2967–E2976. doi:10.1073/pnas.1506763112

Newman ZL, Hoagland A, Aghi K, Worden K, Levy SL, Son JH, Lee LP, Isacoff EY. 2017. Input-Specific Plasticity and Homeostasis at the Drosophila Larval Neuromuscular Junction. Neuron 93:1388–1404.e10. doi:10.1016/j.neuron.2017.02.028

Nguyen TH, Vicidomini R, Choudhury SD, Coon SL, Iben J, Brody T, Serpe M. 2021. Single-Cell RNA Sequencing Analysis of the Drosophila Larval Ventral Cord. Curr Protoc 1:e38. doi:10.1002/cpz1.38

Nose A. 2012. Generation of neuromuscular specificity in Drosophila: novel mechanisms revealed by new technologies. Front Mol Neurosci 5:62. doi:10.3389/fnmol.2012.00062

Nose A, Mahajan VB, Goodman CS. 1992. Connectin: A homophilic cell adhesion molecule expressed on a subset of muscles and the motoneurons that innervate them in Drosophila. Cell 70:553–567. doi:10.1016/0092-8674(92)90426-d

Nose A, Umeda T, Takeichi M. 1997. Neuromuscular target recognition by a homophilic interaction of connectin cell adhesion molecules in Drosophila. Dev Camb Engl 124:1433–41.

Orgogozo V, Grueber WB. 2005. FlyPNS, a database of the Drosophila embryonic and larval peripheral nervous system. Bmc Dev Biol 5:4. doi:10.1186/1471-213x-5-4

Özkan E, Carrillo RA, Eastman CL, Weiszmann R, Waghray D, Johnson KG, Zinn K, Celniker SE, Garcia KC. 2013. An Extracellular Interactome of Immunoglobulin and LRR Proteins Reveals Receptor-Ligand Networks. Cell 154:228–239. doi:10.1016/j.cell.2013.06.006

Pérez-Moreno JJ, O’Kane CJ. 2018. GAL4 Drivers Specific for Type Ib and Type Is Motor Neurons in Drosophila. G3 Genes Genomes Genetics g3.200809.2018. doi:10.1534/g3.118.200809

Ponton F, Chapuis M-P, Pernice M, Sword GA, Simpson SJ. 2011. Evaluation of potential reference genes for reverse transcription-qPCR studies of physiological responses in Drosophila melanogaster. J Insect Physiol 57:840–850. doi:10.1016/j.jinsphys.2011.03.014

Rose D, Zhu X, Kose H, Hoang B, Cho J, Chiba A. 1997. Toll, a muscle cell surface molecule, locally inhibits synaptic initiation of the RP3 motoneuron growth cone in Drosophila. Dev Camb Engl 124:1561–71.

Sanes JR, Zipursky SL. 2020. Synaptic Specificity, Recognition Molecules, and Assembly of Neural Circuits. Cell 181:536–556. doi:10.1016/j.cell.2020.04.008

Saunders HAJ, Johnson-Schlitz DM, Jenkins BV, Volkert PJ, Yang SZ, Wildonger J. 2021. Acetylated α-tubulin residue K394 regulates microtubule stability to shape the growth of axon terminals. BioRxiv. doi:10.1101/2021.04.01.438108

Schmid A, Chiba A, Doe CQ. 1999. Clonal analysis of Drosophila embryonic neuroblasts: neural cell types, axon projections and muscle targets. Development 126:4653–4689. doi:10.1242/dev.126.21.4653

Schrader S, Merritt DJ. 2000. Central projections of Drosophila sensory neurons in the transition from embryo to larva. J Comp Neurol 425:34–44. doi:10.1002/1096-9861(20000911)425:1<34::aid-cne4>3.0.co;2-g

Shen K, Bargmann CI. 2003. The Immunoglobulin Superfamily Protein SYG-1 Determines the Location of Specific Synapses in C. elegans. Cell 112:619–630. doi:10.1016/s0092-8674(03)00113-2

Shen K, Fetter RD, Bargmann CI. 2004. Synaptic Specificity Is Generated by the Synaptic Guidepost Protein SYG-2 and Its Receptor, SYG-1. Cell 116:869–881. doi:10.1016/s0092-8674(04)00251-x

Shishido E, Takeichi M, Nose A. 1998. Drosophila Synapse Formation: Regulation by Transmembrane Protein with Leu-Rich Repeats, CAPRICIOUS. Science 280:2118–2121. doi:10.1126/science.280.5372.2118

Sink H, Whitington P. 1991a. Pathfinding in the central nervous system and periphery by identified embryonic Drosophila motor axons. Development (Cambridge, England) 112:307—316.

Sink H, Whitington PM. 1991b. Location and connectivity of abdominal motoneurons in the embryo and larva of Drosophila melanogaster. J Neurobiol 22:298–311. doi:10.1002/neu.480220309

Soba P, Zhu S, Emoto K, Younger S, Yang S-J, Yu H-H, Lee T, Jan LY, Jan Y-N. 2007. Drosophila Sensory Neurons Require Dscam for Dendritic Self-Avoidance and Proper Dendritic Field Organization. Neuron 54:403–416. doi:10.1016/j.neuron.2007.03.029

Szymczak-Workman AL, Vignali KM, Vignali DAA. 2012. Design and Construction of 2A Peptide-Linked Multicistronic Vectors. Cold Spring Harb Protoc 2012:pdb.ip067876. doi:10.1101/pdb.ip067876

Takizawa E, Komatsu A, Tsujimura H. 2007. Identification of Common Excitatory Motoneurons in Drosophila melanogaster Larvae. Zool Sci 24:504–513. doi:10.2108/zsj.24.504

Tang F, Barbacioru C, Wang Y, Nordman E, Lee C, Xu N, Wang X, Bodeau J, Tuch BB, Siddiqui A, Lao K, Surani MA. 2009. mRNA-Seq whole-transcriptome analysis of a single cell. Nat Methods 6:377–382. doi:10.1038/nmeth.1315

Thor S, Thomas JB. 1997. The Drosophila islet Gene Governs Axon Pathfinding and Neurotransmitter Identity. Neuron 18:397–409. doi:10.1016/s0896-6273(00)81241-6

Valdes-Aleman J, Fetter RD, Sales EC, Heckman EL, Venkatasubramanian L, Doe CQ, Landgraf M, Cardona A, Zlatic M. 2021. Comparative Connectomics Reveals How Partner Identity, Location, and Activity Specify Synaptic Connectivity in Drosophila. Neuron 109. doi:10.1016/j.neuron.2020.10.004

Veling MW, Li Y, Veling MT, Litts C, Michki N, Liu H, Ye B, Cai D. 2019. Identification of Neuronal Lineages in the Drosophila Peripheral Nervous System with a “Digital” Multi-spectral Lineage Tracing System. Cell Reports 29:3303–3312.e3. doi:10.1016/j.celrep.2019.10.124

Venkatasubramanian L, Guo Z, Xu S, Tan L, Xiao Q, Nagarkar-Jaiswal S, Mann RS. 2019. Stereotyped terminal axon branching of leg motor neurons mediated by IgSF proteins DIP-α and Dpr10. Elife 8:e42692. doi:10.7554/elife.42692

Venken KJT, Schulze KL, Haelterman NA, Pan H, He Y, Evans-Holm M, Carlson JW, Levis RW, Spradling AC, Hoskins RA, Bellen HJ. 2011. MiMIC: a highly versatile transposon insertion resource for engineering Drosophila melanogaster genes. Nat Methods 8:737–743. doi:10.1038/nmeth.1662

Vicidomini R, Nguyen TH, Choudhury SD, Brody T, Serpe M. 2021. Assembly and Exploration of a Single Cell Atlas of the Drosophila Larval Ventral Cord. Identification of Rare Cell Types. Curr Protoc 1:e37. doi:10.1002/cpz1.37

Wang J, Ma X, Yang JS, Zheng X, Zugates CT, Lee C-HJ, Lee T. 2004. Transmembrane/Juxtamembrane Domain-Dependent Dscam Distribution and Function during Mushroom Body Neuronal Morphogenesis. Neuron 43:663–672. doi:10.1016/j.neuron.2004.06.033

Wang Y, Lobb-Rabe M, Ashley J, Anand V, Carrillo RA. 2021. Structural and functional synaptic plasticity induced by convergent synapse loss in the Drosophila neuromuscular circuit. J Neurosci JN-RM-1492-20. doi:10.1523/jneurosci.1492-20.2020

Winberg ML, Mitchell KJ, Goodman CS. 1998. Genetic Analysis of the Mechanisms Controlling Target Selection: Complementary and Combinatorial Functions of Netrins, Semaphorins, and IgCAMs. Cell 93:581–591. doi:10.1016/s0092-8674(00)81187-3

Wit J de, Ghosh A. 2016. Specification of synaptic connectivity by cell surface interactions. Nat Rev Neurosci 17:4–4. doi:10.1038/nrn.2015.3

Xie Q, Brbic M, Horns F, Kolluru SS, Jones RC, Li J, Reddy AR, Xie A, Kohani S, Li Z, McLaughlin CN, Li T, Xu C, Vacek D, Luginbuhl DJ, Leskovec J, Quake SR, Luo L, Li H. 2021. Temporal evolution of single-cell transcriptomes of Drosophila olfactory projection neurons. Elife 10:e63450. doi:10.7554/elife.63450

Xu C, Theisen E, Maloney R, Peng J, Santiago I, Yapp C, Werkhoven Z, Rumbaut E, Shum B, Tarnogorska D, Borycz J, Tan L, Courgeon M, Griffin T, Levin R, Meinertzhagen IA, Bivort B de, Drugowitsch J, Pecot MY. 2019. Control of Synaptic Specificity by Establishing a Relative Preference for Synaptic Partners. Neuron 103:865–877.e7. doi:10.1016/j.neuron.2019.06.006

Xu S, Sergeeva AP, Katsamba PS, Mannepalli S, Bahna F, Bimela J, Zipursky SL, Shapiro L, Honig B, Zinn K. 2021. Affinity requirements for control of synaptic targeting and neuronal cell survival by heterophilic IgSF cell adhesion molecules. BioRxiv. doi:10.1101/2021.02.16.431482

Xu S, Xiao Q, Cosmanescu F, Sergeeva AP, Yoo J, Lin Y, Katsamba PS, Ahlsen G, Kaufman J, Linaval NT, Lee P-T, Bellen HJ, Shapiro L, Honig B, Tan L, Zipursky SL. 2018. Interactions between the Ig-Superfamily Proteins DIP-α and Dpr6/10 Regulate Assembly of Neural Circuits. Neuron 100:1369–1384.e6. doi:10.1016/j.neuron.2018.11.001

Yamagata M, Sanes JR. 2019. Expression and Roles of the Immunoglobulin Superfamily Recognition Molecule Sidekick1 in Mouse Retina. Front Mol Neurosci 11:485. doi:10.3389/fnmol.2018.00485

Yildirim K, Petri J, Kottmeier R, Klämbt C. 2019. Drosophila glia: Few cell types and many conserved functions. Glia 67:5–26. doi:10.1002/glia.23459

Yoshihara M, Rheuben MB, Kidokoro Y. 1997. Transition from Growth Cone to Functional Motor Nerve Terminal inDrosophila Embryos. J Neurosci 17:8408–8426. doi:10.1523/jneurosci.17-21-08408.1997

Zarin AA, Labrador J-P. 2019. Motor axon guidance in Drosophila. Semin Cell Dev Biol 85:36–47. doi:10.1016/j.semcdb.2017.11.013

Zarin AA, Mark B, Cardona A, Litwin-Kumar A, Doe CQ. 2019. A multilayer circuit architecture for the generation of distinct locomotor behaviors in Drosophila. Elife 8:e51781. doi:10.7554/elife.51781

Zhan X-L, Clemens JC, Neves G, Hattori D, Flanagan JJ, Hummel T, Vasconcelos ML, Chess A, Zipursky SL. 2004. Analysis of Dscam Diversity in Regulating Axon Guidance in Drosophila Mushroom Bodies. Neuron 43:673–686. doi:10.1016/j.neuron.2004.07.020

Zhang H, Rigo F, Martinson HG. 2015. Poly(A) Signal-Dependent Transcription Termination Occurs through a Conformational Change Mechanism that Does Not Require Cleavage at the Poly(A) Site. Mol Cell 59:437–448. doi:10.1016/j.molcel.2015.06.008

Zinn K, Özkan E. 2017. Neural immunoglobulin superfamily interaction networks. Curr Opin Neurobiol 45:99–105. doi:10.1016/j.conb.2017.05.010

